# Seq-Scope-eXpanded: Spatial Omics Beyond Optical Resolution

**DOI:** 10.1101/2025.02.04.636355

**Authors:** Angelo Anacleto, Weiqiu Cheng, Qianlu Feng, Chun-Seok Cho, Yongha Hwang, Yongsung Kim, Yichen Si, Anna Park, Jer-En Hsu, Mitchell Schrank, Rosane Teles, Robert L. Modlin, Olesya Plazyo, Johann E. Gudjonsson, Myungjin Kim, Chang H. Kim, Hee-Sun Han, Hyun Min Kang, Jun Hee Lee

## Abstract

Sequencing-based spatial transcriptomics (sST) enables transcriptome-wide gene expression mapping but falls short of reaching the optical resolution (200–300 nm) of imaging-based methods. Here, we present Seq-Scope-X (Seq-Scope-eXpanded), which empowers submicrometer-resolution Seq-Scope with tissue expansion to surpass this limitation. By physically enlarging tissues, Seq-Scope-X minimizes transcript diffusion effects and increases spatial feature density by an additional order of magnitude. In liver tissue, this approach resolves nuclear and cytoplasmic compartments in nearly every single cell, uncovering widespread differences between nuclear and cytoplasmic transcriptome patterns. Independently confirmed by imaging-based methods, these results suggest that individual hepatocytes can dynamically switch their metabolic roles. Seq-Scope-X is also applicable to non-hepatic tissues such as brain and colon, and can be modified to perform spatial proteomic analysis, simultaneously profiling hundreds of barcode-tagged antibody stains at microscopic resolutions in mouse spleens and human tonsils. These findings establish Seq-Scope-X as a transformative tool for ultra-high-resolution whole-transcriptome and proteome profiling, offering unparalleled spatial precision and advancing our understanding of cellular architecture, function, and disease mechanisms.

## Introduction

Spatial transcriptomics (ST) has revolutionized our ability to interrogate gene expression within intact tissues, enabling studies of cellular organization, function, and interactions in situ [1]. By mapping the transcriptome directly onto tissue sections, ST reveals intricate spatial patterns critical for understanding complex processes such as development, homeostasis, and disease pathology [1–5].

Among various ST techniques, sequencing-based approaches (sST) [4] combine spatially coordinated nucleotide barcodes with Next-Generation Sequencing (NGS) to resolve transcript locations. These techniques, including the original ST [6], 10x Visium [7], Slide-Seq [8, 9], DBiT-seq [10], and HDST [11], offer genome-wide coverage and have progressively achieved higher spatial resolution over time. However, even the latest sST platforms—such as Seq-Scope [12], Stereo-Seq [13], Pixel-Seq [14], as well as Seq-Scope’s recent variations (e.g., including Open-ST and Nova-ST) [15–17]—struggle to reach the resolution of optical microscopy (∼200 nm). The spatial barcode spacing is inherently constrained to ∼0.5 µm by the procedures underlying NGS-based spatial barcode sequencing. Additionally, RNA diffusion during tissue permeabilization and capture further exacerbates this limitation, reducing the effective resolution to approximately 1–3 µm [1–3].

In contrast, imaging-based ST (iST) [4] methods—such as MERFISH [18], CosMx [19], and Xenium [20]—achieve resolutions as fine as ∼200 nm by optically detecting fluorescently labeled RNAs through sequential hybridization or in situ sequencing. Tissue expansion techniques have further advanced iST resolution into the nanoscale regime [21, 22]. However, iST methods are generally constrained to analyzing a predefined gene set, limiting their ability to comprehensively capture global transcriptomic complexity, including splicing variants and somatic mutations. Scaling iST methods to encompass global transcriptomic complexity becomes prohibitively expensive due to the need for unique imaging probes, which significantly increases cost. Moreover, while sST methods are time-efficient, completing experiments in a single measurement, iST methods are time-intensive, requiring repeated imaging– erasing cycles to detect multiple targets. These iterative cycles not only limit scalability and practicality [4, 5], but also introduce cumulative errors—such as photobleaching, incomplete probe removal, and imaging artifacts—that compromise precision and diminish the resolution advantage [4, 5]. Consequently, a critical gap persists: no existing platform currently offers whole-transcriptome coverage at a reasonable time and cost while achieving spatial resolution at or beyond the diffraction limit of optical microscopy.

Here, we introduce Seq-Scope-X (Seq-Scope-eXpanded), a platform that delivers super-resolution spatial analysis—surpassing the diffraction limit of optical microscopy—while preserving the whole-transcriptome coverage and cost-effectiveness of the original Seq-Scope. Inspired by tissue expansion– based approaches in iST [21, 22] and their integration into lower-resolution sST workflows [23], Seq-Scope-X synergistically incorporates tissue expansion into Seq-Scope’s submicrometer-resolution capabilities. This innovation increases spatial feature density by an additional order of magnitude, generating tens of millions of hexagonally arranged coordinate points per square millimeter. Furthermore, proportional tissue expansion reduces distortion and limits RNA diffusion, enhancing spatial precision beyond what was achieved by existing variations of Seq-Scope [15–17].

In addition to providing unprecedented resolution, Seq-Scope-X facilitates high-resolution multi-omics profiling by decoupling molecular probe staining from spatial capture. We demonstrate that Seq-Scope-X supports massively multiplexed proteomics analysis using hundreds of barcode-tagged antibodies, while maintaining the same resolution and throughput as its transcriptomic assays. By bridging the gap between comprehensive NGS-based profiling and the nanoscale precision of imaging-based methods, this integrated approach paves the way for unparalleled insights into cellular architecture, subcellular heterogeneity, and disease mechanisms across diverse tissues.

## Results

### Seq-Scope-X Workflow

The Seq-Scope-X protocol builds on two recent advances in spatial transcriptomics: the tissue expansion method for enhanced resolution [24] and the submicrometer-resolution Seq-Scope platform [12, 15]. By integrating these approaches, Seq-Scope-X achieves sub-200 nm super-resolution spatial transcriptomics, surpassing the diffraction limit of optical microscopy. Frozen tissue sections were fixed (Fig. 1A), permeabilized, and hybridized with locked nucleic acid (LNA)-containing oligo-dT probes labeled with a red fluorescent dye and an acrydite motif (Fig. 1B). The hybridized tissue was embedded in an expandable polyacrylamide gel, incorporating the acrydite-bound oligo-dT and associated mRNA species into the polymer matrix (Fig. 1C). After embedding, the tissue was digested and cleared with proteinase K, and the gel was expanded by placing it in a hypotonic buffer. The expanded gel, now containing spatially preserved mRNAs, was overlaid onto the Seq-Scope’s spatially barcoded array (Chip), which was produced using a NovaSeq 6000 S4 flow cell [15]. mRNA transfer from the gel to the Seq-Scope Chip was facilitated by heating the gel-array sandwich at 45°C (Fig. 1D). Finally, the Seq-Scope Chip, with mRNAs transferred from the gel, was processed into spatially labeled cDNA libraries using the conventional Seq-Scope library prep method [12, 15].

**Fig 1.**
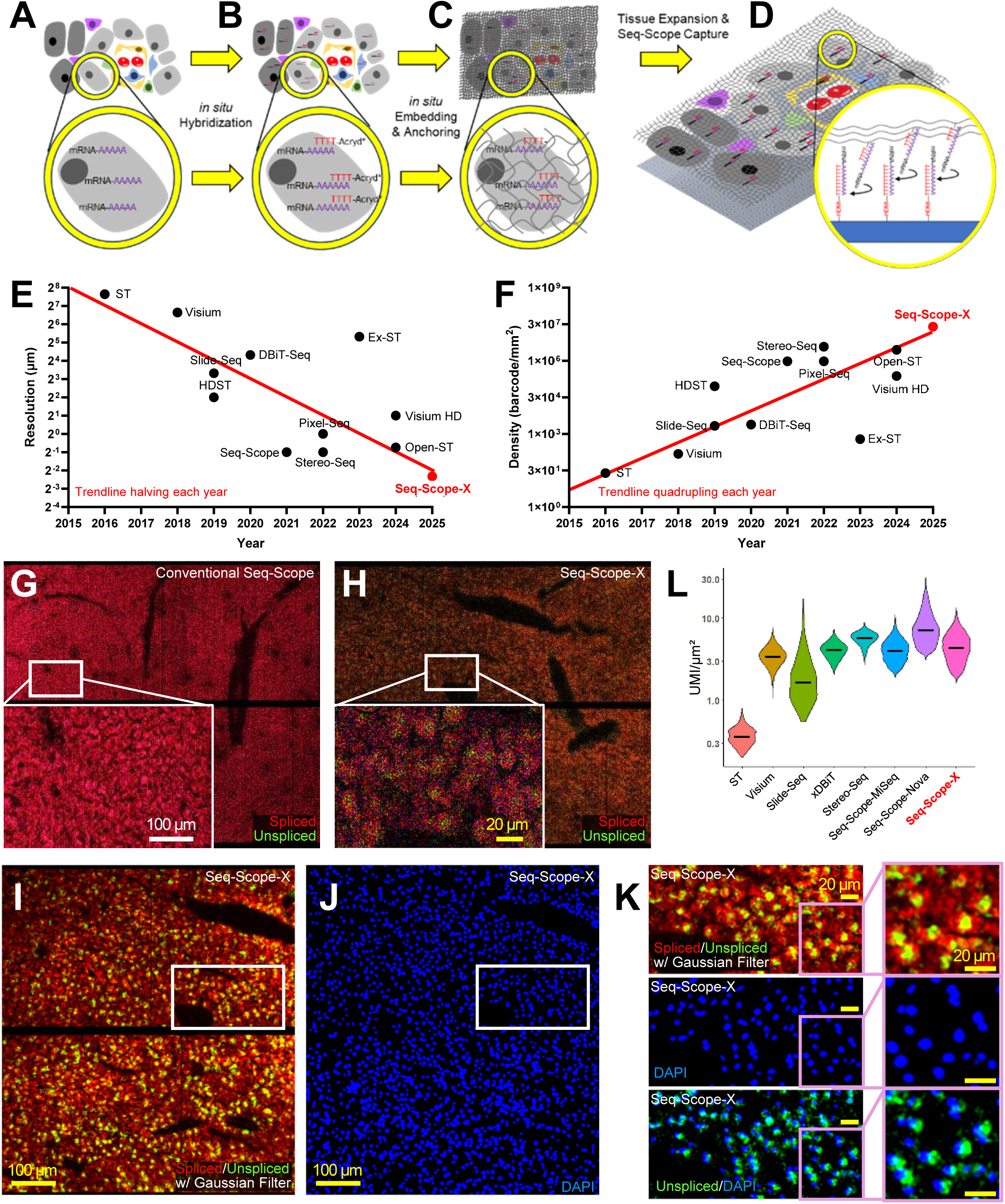
Methodology and Performance of Seq-Scope-X. (A-D) Schematic diagram depicting the Seq-Scope-X procedure. Tissue (A) was first hybridized with fluorescence-labeled (*, ATTO590) oligo-dT acrydite probe (B). Then it was embedded in an acrylite hydrogel (C), which was expanded after tissue digestion. (D) The mRNAs were then transferred to the Seq-Scope array through a heat-mediated template switching process for a spatial transcriptomics analysis. (E and F) Comparison of different spatial transcriptomics (ST) technologies based on their spatial performance. The scatterplots depict the X-axis as the release date (commercial or other unpublished technologies) or peer-reviewed publication date (published technologies), and the Y-axis as the resolution (E) and spatial pixel density throughput (F). A red trendline illustrates the improvement trends: resolution doubling approximately every year (E), and spatial pixel density increasing approximately fourfold each year (F). (G and H) Spatial plotting of spliced (red) and unspliced (green) transcripts in conventional Seq-Scope (G) and Seq-Scope-X (H) data. (I-K) Digitally enhanced images of spliced (red) and unspliced (green) transcripts processed using Gaussian filtering (I and K) to reflect the probabilistic local diffusion of transcripts. These images are overlaid with a DAPI-stained image (J and K), which is spatially aligned with the transcriptomic data. (L) Comparison of different ST technologies on their transcript capture efficiency. Seq-Scope-X performance is comparable to most state-of-the-art ST technologies having relatively lower resolutions. White scale bars represent the original scale, while yellow scale bars indicate the scale adjusted to account for tissue expansion.

### Seq-Scope-X Tissue Expansion Maintains Structural Integrity and Uniformity

We first verified that tissue expansion in our workflow could be achieved without substantial distortion of the overall tissue structure, a critical requirement for accurate spatial transcriptomics. The expansion scale factor consistently ranged between 2-and 3-fold across experimental batches and was easily determined by measuring the gel size before and after expansion. DAPI-labeled DNA remained stably anchored within the gel, indicating minimal diffusion of nuclear content, while the tissue expanded uniformly across the entire section, preserving the relative positions and proportional areas of nuclei (Fig. S1A, S1B). Computational dilation of images further demonstrated that nuclear positions before and after expansion were nearly identical across most tissue regions (Fig. S1A, S1B). Additionally, red fluorescence signals from the oligo-dT-acrydite probe remained localized around the DAPI-labeled nuclei after tissue expansion (Fig. S1C), consistent with the high concentration of mRNA in the perinuclear region where translation actively occurs.

### Seq-Scope-X Achieves Sub-200 nm Supra-Optical Resolution

The latest version of Seq-Scope and its variations achieve a resolution of 0.6 µm in a hexagonally arranged pixel array [15–17], with a pixel density of approximately 3 million per mm². However, when combined with the 3-fold tissue expansion procedure in Seq-Scope-X, the resolution is effectively improved to 0.2 µm and the pixel density is increased to approximately 27 million per mm². This represents a substantial enhancement over existing sequencing-based spatial transcriptomics (sST) technologies in both resolution (Fig. 1E) and spatial feature density (Fig. 1F), enabling a much more detailed view of the tissue transcriptome, particularly in densely packed or subcellular regions.

To evaluate whether Seq-Scope-X maintains its high-resolution utility in practical tissue analyses, we applied it to liver tissue—a model well-suited for cross-dataset comparisons due to its repeating parenchymal units and well-defined metabolic zonation layer structure. Liver tissue minimizes common challenges in benchmarking procedures such as variability in tissue depth, orientation, and cell type composition. Data from Seq-Scope-X were directly compared with those from the original Seq-Scope, confirming that Seq-Scope-X produced the anticipated spatial readout with substantially enhanced resolution and throughput. While the original Seq-Scope could map spliced (red) and unspliced (green) transcripts to discern single-cell hepatocellular structures and occasional subcellular features, single-cell boundaries were often diffuse, and subcellular details were ambiguous in most cases (Fig. 1G). In contrast, Seq-Scope-X distinctly resolved single cells with clear extracellular domains lacking transcripts, while also identifying unspliced RNAs with strong nuclear-like localization at the center of hepatocyte regions (Fig. 1H).

Simulating Gaussian RNA diffusion with a maximum range of 1 µm (scaled to ∼330 nm by the expansion factor, with most reads concentrated within a ∼100 nm radius) allowed Seq-Scope-X to more precisely identify nuclei positions as clusters of unspliced RNA (Fig. 1I). The unspliced transcript clusters were highly consistent with DAPI-stained nuclear regions (Fig. 1J), underscoring Seq-Scope-X’s microscopic precision (Fig. 1K). These findings confirm that Seq-Scope-X achieves its advanced resolution in experimental applications, enabling the visualization of subcellular transcriptomic structures across the tissue section.

### Seq-Scope-X Retains High RNA Capture Efficiency

While Seq-Scope-X’s resolution was clearly enhanced, another important consideration is whether it provides sufficient transcriptome coverage comparable to that of the original Seq-Scope and other ST methodologies. To address this, we analyzed seven publicly available ST datasets, all prepared from C57BL/6 WT mouse liver, including those from the original ST (Fig. S1D) [25], 10x Visium (Fig. S1E) [26], Slide-Seq (Fig. S1F) [8], xDBiT (an improved version of DBiT-Seq; Fig. S1G) [27], Seq-Scope^MISEQ^ (original Seq-Scope implemented with Illumina MISEQ; Fig. S1H) [12], Stereo-Seq (Fig. S1I) [28], and Seq-Scope^NOVASEQ^ (recently updated Seq-Scope implemented with Illumina NOVASEQ 6000 [15], similar to Open-ST [16] and Nova-ST [17]; Fig. S1J), alongside the Seq-Scope-X data (Fig. S1K). In an area-normalized comparison of transcript capture efficiency, Seq-Scope-X demonstrated excellent performance per µm² (Fig. 1L), surpassing most datasets except Stereo-Seq and Seq-Scope^NOVASEQ^, which showed comparable performance in this analysis (Fig. 1L, S1D-S1K). These results underscore the capability of Seq-Scope-X to achieve high transcript capture efficiency compared to existing approaches.

### Seq-Scope-X Matches Bulk RNA-seq Accuracy

The transcriptome content of Seq-Scope-X was highly similar to bulk RNA-seq [29], conventional Seq-Scope [12, 15], and other ST techniques [8, 26–28], showing the correlation coefficient around 0.9 in most comparisons (Fig. S2A, S2B). These results demonstrate that, despite its multi-step process (Fig. 1A), Seq-Scope-X maintains transcriptome accuracy comparable to both its predecessor and other spatial transcriptomics methods.

### Seq-Scope-X Reveals Nuclear-Cytoplasmic Transcriptome Disparities

Using Seq-Scope-X, we performed subcellular-level analysis by leveraging the distribution of spliced and unspliced transcripts, which represent cell boundary and nuclear positions, respectively (Fig. 1I–K).

Using this image, the Watershed cell segmentation algorithm divided the spatial area into single cells and labeled the local maxima of unspliced transcripts in each segment. This information was then used to define nuclei and cytoplasm (Fig. 2A). Each nuclear and cytoplasmic area pair constituted a single cell area (Fig. 2B), and was subjected to a cell type clustering analysis (Fig. 2C, 2D).

**Fig 2.**
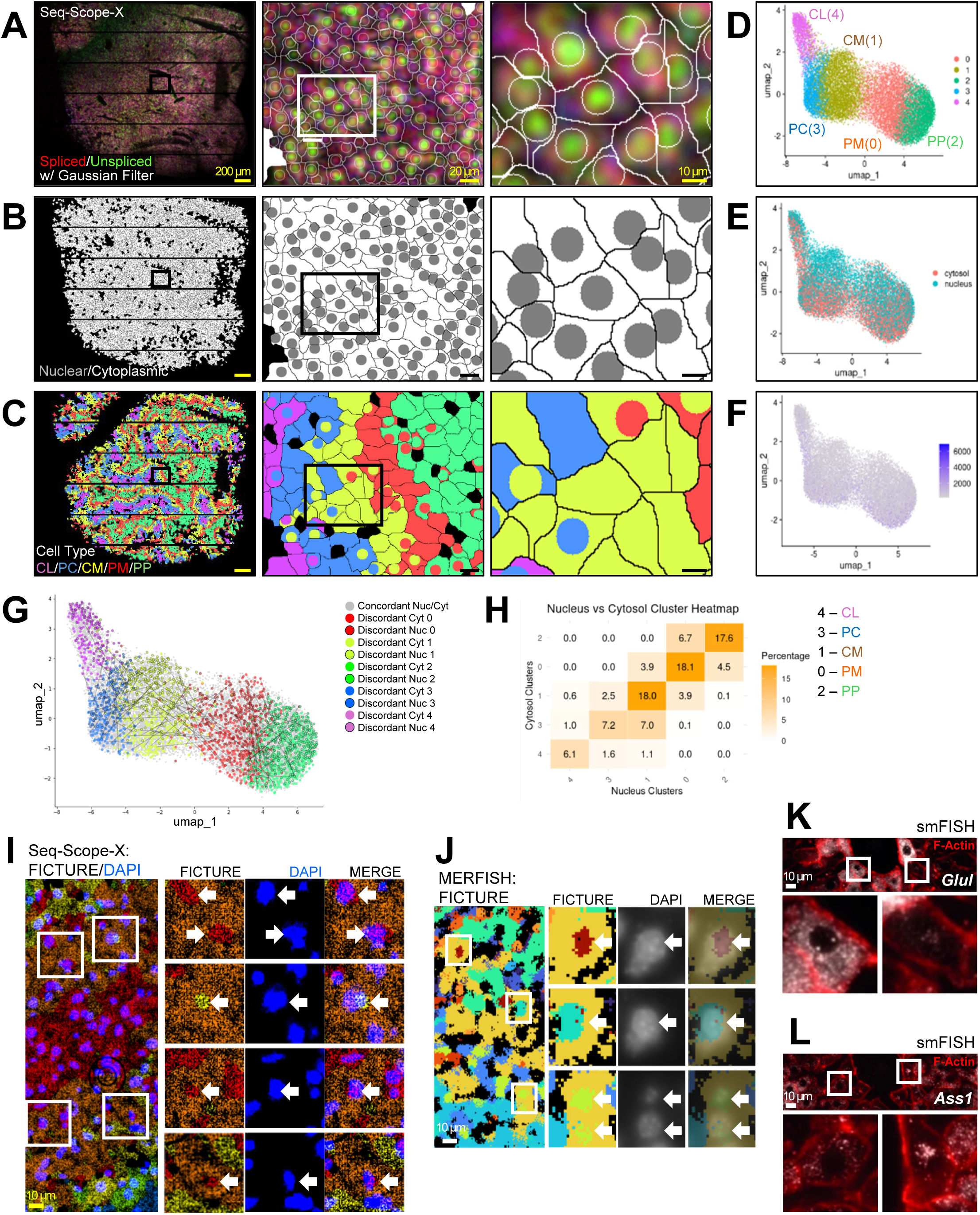
Seq-Scope-X Reveals Subcellular Heterogeneity in Liver Spatial Transcriptome (A-H) Segmentation-based nuclear-cytoplasmic spatial fractionation analysis. (A) Gaussian-filtered unspliced (green)/spliced (red) gene expression data segmented into single nuclei and cytoplasmic areas using the Watershed algorithm. Segment boundaries are indicated in white. (B) Nuclear segments are shown in grey, and cytoplasmic segments in white. (C) Segments are color-coded based on cell type mapping results, with the color scheme used in (D). The five cell types correspond to centrilobular (CL; magenta), pericentral (PC; blue), central-side midzone (CM; yellow), portal-size midzone (PM; red), and periportal (PP; green) hepatocytes. In (A-C), boxed regions in the leftmost image are magnified in the center, and the boxed regions in the center image are further magnified on the right. (D-F) UMAP visualization of cell type clustering based on nuclear and cytoplasmic segments. (D) Cell type clusters are color-coded, matching the scheme in (C). (E) Each data point is color-coded to indicate nuclear (green) or cytoplasmic (red) segments. (F) Each data point is colored by the number of unique transcripts identified in each segment. (G) UMAP manifold from (D) highlights nuclear-cytoplasmic pairs from single cells, connected by lines to show relationships between nuclear and cytoplasmic transcriptomes. The analysis reveals widespread mismatches between nuclear and cytoplasmic transcriptome phenotypes. (H) Heatmap (H) illustrating mismatches between nuclear and cytoplasmic cell types within neighboring hepatocellular zonation clusters (4-3-1-0-2; CL-PC-CM-PM-PP). (I) Segmentation-free FICTURE analysis of hepatocellular cell types from Seq-Scope-X data (as shown in Fig. 3) reveals a pixel-level cell type factor map, matched with DAPI (blue) images. Boxed areas are magnified in right. Arrows highlight nuclear-cytoplasmic mismatches in cell type factors. (J) Segmentation-free FICTURE analysis of hepatocellular cell types from MERSCOPE data [31] reveals a pixel-level cell type factor map. Magnified images (second column) are matched (third column) and merged (fourth column) with DAPI (white) images. Nuclear-cytoplasmic mismatches are highlighted by arrows. (K, L) Single-molecule RNA in situ hybridization (smFISH) imaging of the indicated genes (white). Cell boundaries are visualized through F-actin staining (red). Magnified white-boxed regions highlight cells with cytoplasmic (lower left) or nuclear (lower right) gene expression, confirming the subcellular transcriptome heterogeneity observed in Seq-Scope-X data. Reproduced with permission from © 2017 Springer Nature [32]. All rights reserved. White scale bars represent the original scale, while yellow and black scale bars indicate the scale adjusted to account for tissue expansion.

Both nuclear and cytoplasmic transcriptomes clustered into five major hepatocellular types, which correspond to periportal (PP), portal-side midzone (PM), central-side midzone (CM), pericentral (PC), and centrilobular (CL) hepatocytes (Fig. 2D). While nuclear and cytoplasmic transcriptomes integrated well within each hepatocellular cluster (Fig. 2E), they displayed notable differences in nuclear-and mitochondrial-specific transcripts (Fig. S2C), represented as a uniform shift in the UMAP manifold (Fig. 2E). Additionally, cytoplasmic clusters generally exhibited a higher number of unique transcripts compared to nuclear clusters (Fig. 2F, S2D).

In the histological space, five hepatocellular types were arranged nicely corresponding to the portal-central axis of liver metabolic zonation, forming a clearly layered structure (Fig. 2C, S2E). Interestingly, while nuclear and cytoplasmic clusters were concordant in most cases, about one third of cells exhibited differing transcriptomic phenotypes between the subcellular compartments (Fig. 2C, S2E; magnified images). These disparities were predominantly observed between clusters of two neighboring zones (Fig. 2C, 2G, 2H, S2E), which represented relatively similar clusters.

In this analysis, the cytoplasmic transcriptome likely reflects a cell’s current phenotype and functional state, while the nuclear transcriptome may indicate its future trajectory in phenotype and function. These findings suggest that hepatocellular zonation and the associated division of labor are dynamic processes, capable of evolving over time or adapting to physiological conditions (e.g., feeding or activity). Although collected during the daytime—a period generally associated with metabolic stability in mice—our data revealed significant fluctuations in hepatocellular phenotypes, highlighting ongoing variability and heterogeneity within and across subcellular compartments.

### Segmentation-Free Analysis Aligns With Nuclear-Cytoplasmic Transcriptome Differences

To validate that our observed subcellular differences were not artifacts of imperfect segmentation or contamination from neighboring cells, we employed FICTURE, a segmentation-free, pixel-level cell type inference method [30]. Using FICTURE, we projected five hepatocellular clusters (PP, PM, CM, PC, and CL) learned from our hexagonal grid analysis (Fig. 3A) into pixel-level histological space. The analysis revealed a clearly layered structure along the portal-central axis (Fig. 3B), though with rough boundaries and scattered fragments between layers (Fig. 3C). When we overlaid the FICTURE factor map with DAPI images, we found that these fragmented clusters aligned closely with nuclear regions where the transcriptome profile differed from that of the surrounding cytoplasm (Fig. 2I). Both our subcellular segmentation analysis (Fig. 2A-2H) and segmentation-free FICTURE analysis (Fig. 2I) of the Seq-Scope-X dataset congruently confirm the presence of subcellular transcriptome heterogeneity, specifically demonstrating distinct nuclear and cytoplasmic transcriptome characteristics within individual cells.

**Fig 3.**
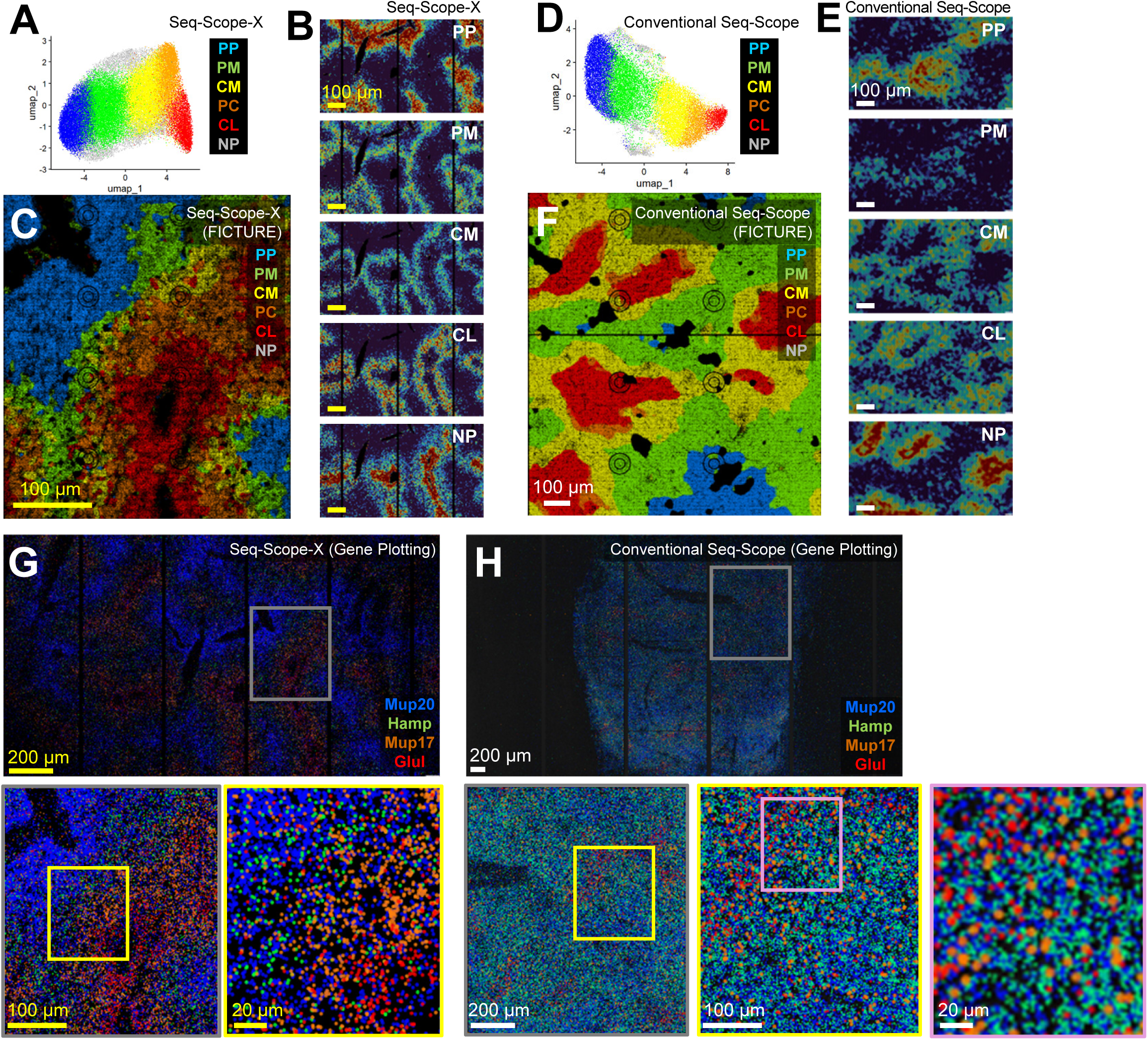
Seq-Scope-X Demonstrates Spatial Precision in Gene Expression and Cell Type Inference. (A and D) Cell type clustering results from Seq-Scope-X (A) and conventional Seq-Scope (D) datasets. Both datasets were processed through 14 μm-sided hexagons (corresponding to the array scale, not accounting for expansion). Both results identified cell types corresponding to Periportal (PP), Portal-and Central-Midzones (PM and CM), Pericentral (PC), Centrilobular (CL) hepatocytes, and non-parenchymal (NP) cell types. The results are presented in UMAP manifolds. For analyses using additional hexagonal sizes, refer to Fig. S3E and S3F. (B, C and E-H) Spatial plot of cell type factors (B, C, E and F) and individual gene expressions (G and H) using the Seq-Scope-X (B, C and G) and conventional Seq-Scope (E, F and H) datasets. The spatial cell type factor maps (B, C, E and F) were constructed using cell type clusters (A and D) through segmentation-free FICTURE projection. Individual factors were projected in a spatial heat map indicating the probability of each factor (B and E). Individual dots in the spatial gene expression plot (G and H) represent the digital gene expression of indicated genes in the same color. White scale bars represent the original scale, while yellow scale bars indicate the scale adjusted to account for tissue expansion.

### In Situ Imaging Confirms Subcellular Transcriptome Heterogeneity

To independently validate the observed subcellular transcriptome heterogeneity, particularly the disparity between nuclear and cytoplasmic phenotypes, we analyzed publicly available datasets from imaging-based spatial transcriptomics (iST) techniques with comparable resolution to Seq-Scope-X (∼200 nm).

Using MERFISH mouse liver data, an iST dataset with an optical resolution (200-300 nm) [31], we applied segmentation-free FICTURE analysis [30] to evaluate whether this method could detect similar disparities. Consistent with our Seq-Scope-X findings, MERFISH data revealed distinct hepatocellular phenotypes of metabolic zonation between nuclear and cytoplasmic compartments within the same cell, again observed widely across the dataset (Fig. 2J).

To further assess subcellular differences, we re-examined single-molecule RNA fluorescence in situ hybridization (smFISH) data reported in the literature [32]. Markers for the central-most CM (*Glul*, Fig. 2K) and portal-most PP (*Ass1*, Fig. 2L) demonstrated distinct subcellular transcript localization in single cells. In many cases, transcripts were observed exclusively in either the nucleus (lower left panels) or the cytoplasm (lower right panels), confirming pronounced subcellular differences at the single-transcript level. These findings not only corroborate the observations from Seq-Scope-X but also suggest that hepatocellular phenotypes could dynamically change, with nuclear and cytoplasmic transcriptomes reflecting different functional states. This underscores the robustness of subcellular transcriptome heterogeneity revealed through Seq-Scope-X and its relevance across diverse orthogonal techniques.

### Seq-Scope-X Delivers Spatial Precision for Cell-and Tissue-level Analysis

While conventional Seq-Scope[12] and similar technologies such as Stereo-Seq, Open-ST, and Nova-ST [13, 16, 17], could occasionally distinguish nuclear-specific transcripts in a limited number of cells, the detailed subcellular-level analysis achieved with Seq-Scope-X was impossible using these conventional methods. Using Seq-Scope-X, we were able to identify nearly every nucleus as a spatial cluster of unspliced transcripts, which is surrounded by spliced transcripts of each cell (Fig. S3A). In contrast, conventional Seq-Scope, even at the same magnification, only detected nuclear-cytoplasmic structures in large cells occasionally (Fig. S3B, center; highlighted by arrows), with most cells showing no discernible structures (Fig. S3B, left and right). Gaussian diffusion modeling of transcripts further emphasized these differences: conventional Seq-Scope could not reliably determine nuclear centers in most cells (Fig. S3D), whereas Seq-Scope-X precisely identified nuclear positions in individual cells (Fig. 1I-1K, 2A, S3C). This improved performance appears to stem from both increased resolution and tissue expansion, with the latter reducing effective transcript diffusion distance by the scale factor.

Even for cell-level and tissue-level analysis, Seq-Scope-X provided much higher precision over conventional Seq-Scope. Even though Seq-Scope-X and conventional Seq-Scope data both were able to reveal five major hepatocellular clusters and non-parenchymal cell clusters in hexagonal binning analysis (Fig. 3A, 3D), and generally produced similar clustering patterns with similar size of hexagonal bins (Fig. S3E, S3F), FICTURE-mediated projection of these cell types into pixel-level histological space produced far clearer qualitative differences in spatial details (Fig. 3B, 3C, 3E, 3F). Although Seq-Scope-X data showed clearly layered structures of spatial factors with defined domains (Fig. 3B, 3C), conventional Seq-Scope data showed poorly defined spatial relationships between hepatocellular types, with overlapping or discontinuous layer structures (Fig. 3E, 3F). Subcellular level details shown in Seq-Scope-X data (Fig. 2I, 3C) was also not attainable by conventional Seq-Scope data (Fig. 3F). Gene plotting analysis also clearly showed distinct domains of cluster-specific marker expression in Seq-Scope-X data (Fig. 3G), while conventional Seq-Scope data failed to reveal that and rather showed a continuously diffused pattern (Fig. 3H). These findings underscore Seq-Scope-X’s superior resolution and spatial precision, enabling a detailed and accurate understanding of tissue architecture.

### Seq-Scope-X Performs High-Resolution Spatial Mapping of Brain and Colon Transcriptomes

To evaluate the broad applicability of Seq-Scope-X, we analyzed brain and colon tissues, which are frequently studied using ST technologies. In brain tissue, Seq-Scope-X effectively captured cellular heterogeneity while maintaining high spatial resolution. The spatial distribution of spliced and unspliced transcripts aligned well with DAPI imaging from the same section (Fig. 4A), validating the spatial precision and subcellular capability of the technique. Cell type clustering successfully identified multiple neuronal populations, including inhibitory GABAergic neurons (*Gad1*, *Gad2*, *Gabra1*, *Sst*), excitatory neurons (*Camk2a*, *Ppp3ca*, *Slc1a2*), and glutamatergic neurons (*Slc17a6*, *Rora*, *Zic1*; Fig. S4A, S4B).

**Fig 4.**
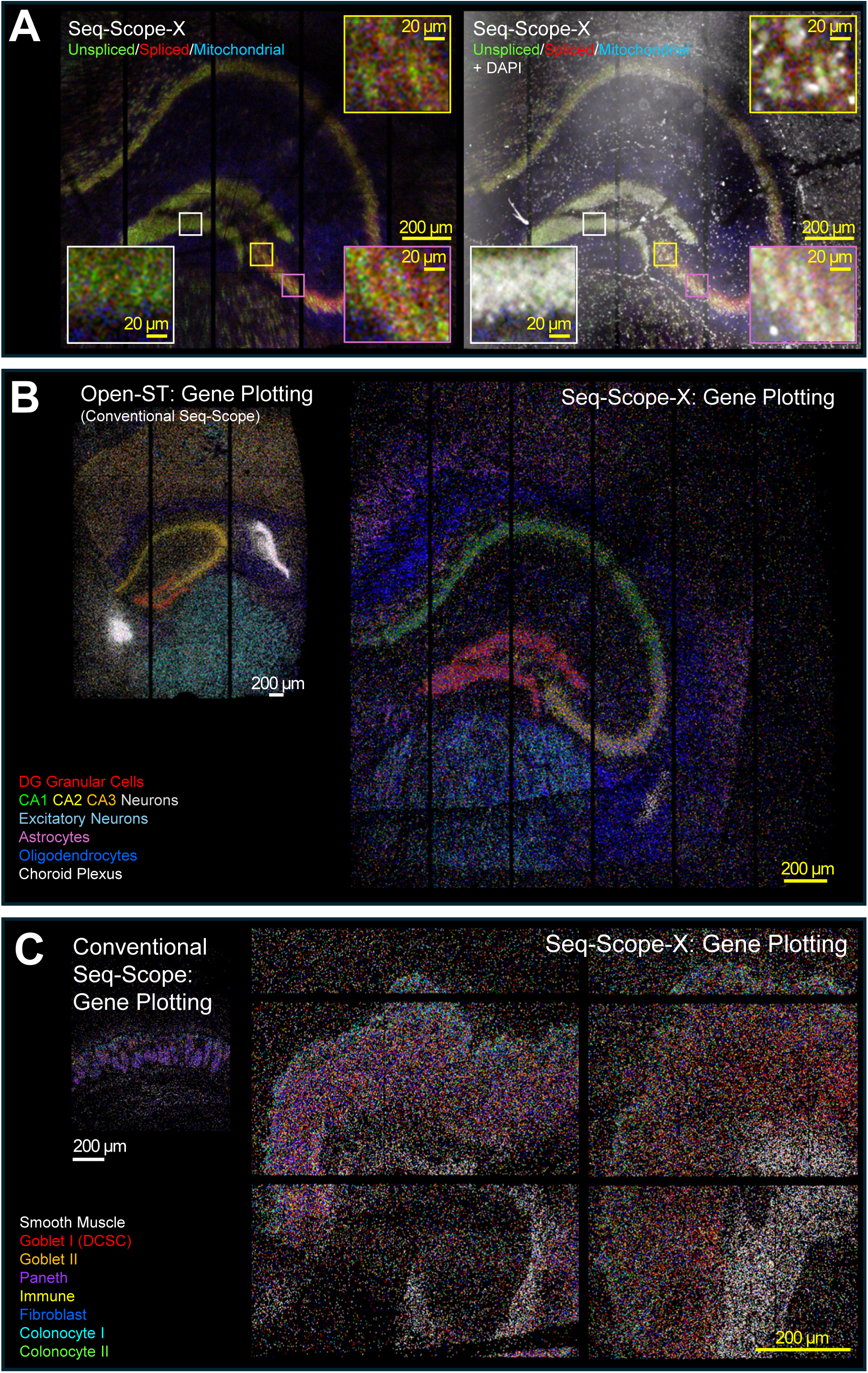
High-Resolution Seq-Scope-X Analysis of Mouse Brain and Colon Tissues. (A) Spatial plotting of spliced (red), unspliced (green), and mitochondrial (blue) transcripts, processed through Gaussian filter, in mouse brain tissue obtained using Seq-Scope-X, shown with DAPI fluorescence image (white, right) of the same expanded tissue section. Insets show magnified views of the regions indicated by colored boxes. (B and C) Spatial gene expression plots comparing cell type-or region-specific gene expression in conventional Seq-Scope (left) and Seq-Scope-X (right) datasets. Mouse brain samples are shown in (B), and colon samples in (C). For a more detailed analysis of the Seq-Scope-X data, refer to Fig. S4. For the list of genes used for the plotting, see the Methods section. White scale bars represent the original scale, while yellow scale bars indicate the scale adjusted to account for tissue expansion.

Additional populations such as *Foxp2*⁺ neurons, granular cells, pyramidal neurons, and non-neuronal cells (astrocytes, oligodendrocytes, choroid plexus cells) were also clearly distinguished. Segmentation-free FICTURE analysis provided high-resolution spatial mapping of these cell types (Fig. S4C, S4D).

Similarly, Seq-Scope-X analysis of colon tissue revealed detailed cellular organization along the surface-crypt axis (Fig. S4E-S4I). The technique distinguished between crypt-localized (type I, deep crypt secretory cells or DCSC) and surface-localized (type II) goblet cell populations, marked by differential expression of *Zg16*/*Fcgbp* and *Muc2*/*Cryab*/*Agr2* respectively. Other identified populations included colonocytes, Paneth/stem cells, immune cells, smooth muscle cells, fibroblasts, and macrophages.

Nuclear transcriptome signatures characterized by *Malat1* and *Neat1* expression were also detected (Fig. S4F), again confirming the capabilities of detecting subcellular transcriptome structures.

Side-by-side comparison of cell type-and region-specific markers between Seq-Scope-X and conventional Seq-Scope demonstrated that Seq-Scope-X data are consistent with previous datasets while enabling massively magnified spatial analysis (Fig. 4B, 4C). These results establish Seq-Scope-X as a reliable tool for high-resolution spatial transcriptomics across diverse tissue types.

### Seq-Scope-X Profiles Mouse Splenic Proteome using Barcode-Tagged Antibodies

Previously, we attempted to capture spatial proteomic information in conventional Seq-Scope using poly-A–labeled barcoded antibodies, a technique originally developed for single-cell CITE-seq [33, 34] and recently implemented in several low-resolution bST platforms [35–37]. The workflow involved attaching tissue to the Seq-Scope Chip, performing barcoded antibody staining, and then releasing the antibody-attached barcode oligonucleotides for poly-A–based capture. However, despite extensive optimization efforts, generating high-resolution proteomic maps proved difficult due to multiple technical challenges. The primary obstacle was achieving optimal tissue permeabilization - we struggled to establish conditions that would release antibody-tagged barcode sequences while maintaining spatial tissue integrity.

Moreover, the method suffered from high background noise and non-specific signals, likely caused by premature capture of antibody-tagged poly-A tails by the Seq-Scope Chip during the staining process.

Given Seq-Scope-X’s success in transcriptomic applications, we hypothesized it could overcome the proteomic challenges faced by conventional Seq-Scope. The key advantages of Seq-Scope-X were twofold: it separated the antibody staining step from poly-A capture, and it eliminated the need for optimizing tissue permeabilization by stably anchoring poly-A to expansion hydrogel during tissue digestion. To test this approach, we used a mouse spleen tissue and a panel of 119 antibodies targeting distinct mouse cell-surface antigens associated with various immune lineages. Each antibody was labeled with an oligonucleotide containing a unique antibody-derived barcode tag (ADT) and a poly(dA) tail [38]. After staining mouse spleen tissue sections with this antibody cocktail, we subjected the stained slides to the Seq-Scope-X procedure (Fig. 5A).

**Fig 5.**
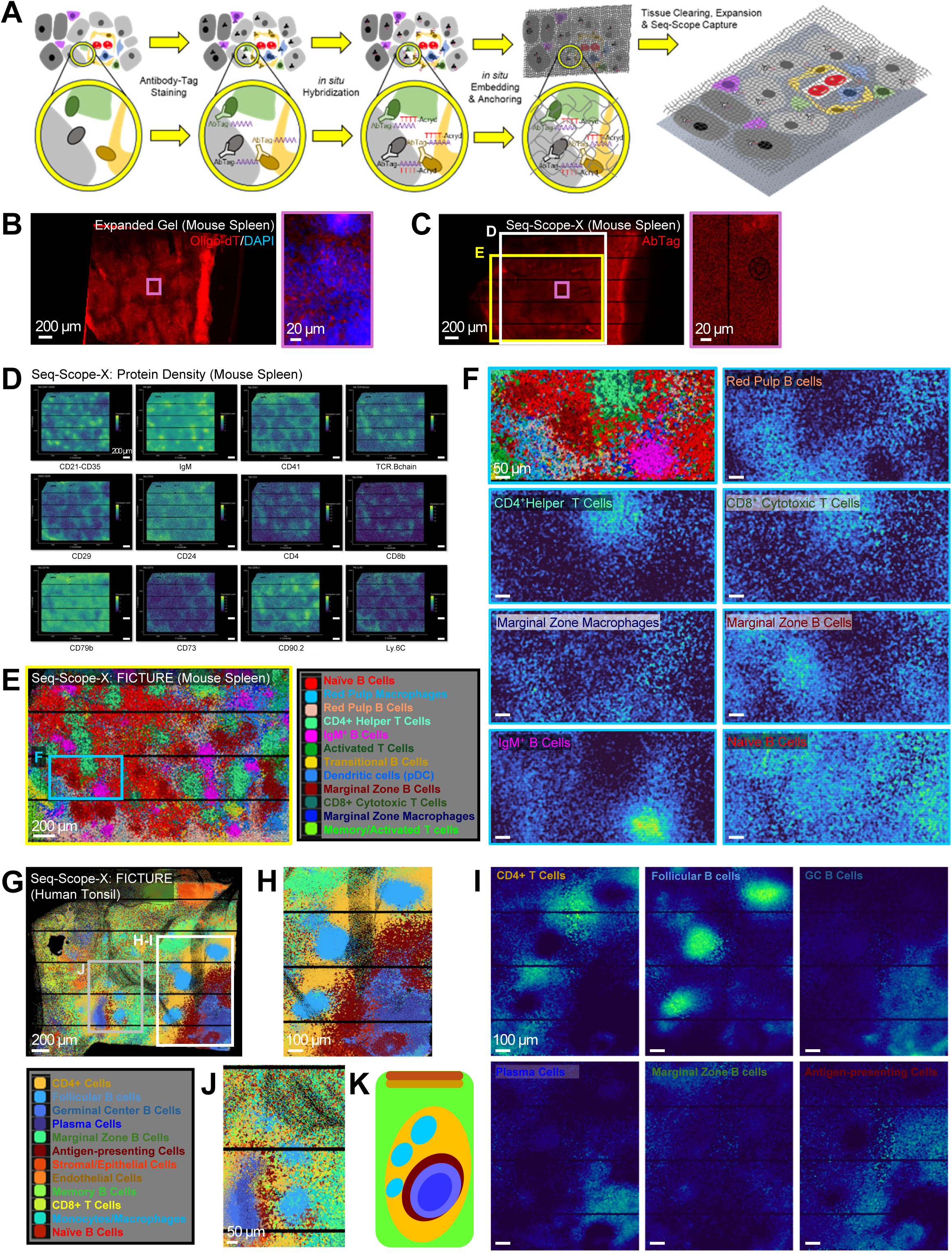
High-Resolution Profiling of Mouse Spleen and Human Tonsil Proteome Using Seq-Scope-X. (A) Schematic of the Seq-Scope-X procedure modified for proteome analysis. Tissues were stained with oligonucleotide barcode-tagged antibodies (antibody tags or AbTags) before undergoing the standard Seq-Scope-X procedure described in Fig. 1A. (B-F) Seq-Scope-X analysis of mouse spleen stained with an AbTag cocktail targeting 119 unique cell surface antigens, including principal lineage markers, plus 9 isotype controls (total 128 AbTags). (B) Fluorescence images of DAPI (blue) and oligo-dT probe (red) post-tissue expansion. The magnified pink boxed area reveals preserved nuclear-membrane structures. The antibody cocktail primarily targeted immune cell surface lineage markers, with signals localized to cell membranes and cell-cell boundaries, distinct from DAPI. The image spatially corresponds to the digitally reconstructed Seq-Scope-X image shown in (C). (C) Spatial plot of antibody tags (red) detected in Seq-Scope-X data. The magnified pink boxed area demonstrates structural similarities with (B) but appears more diffused. Boxed areas in white, yellow, and pink are further analyzed in (D), (E) and (C, right panel), respectively. (D) Representative spatial expression plot of antibody tags targeting specific proteins within the white boxed region in (C). For spatial expression plots of all 128 antibody tags, refer to Fig. S5A. (E and F) FICTURE projection of spatial factors identified via Latent Dirichlet Allocation (LDA) analysis of protein expression results. Twelve distinct factors are annotated and color-coded. The yellow box in (C) is magnified in (E). The blue box in (E) is further magnified in (F). In (E), the positional probability of selected LDA factors was presented in spatial heat maps. For spatial heat maps of all 12 LDA factors for a wider area, refer to Fig. S5B. (G-K) Seq-Scope-X analysis of human tonsil stained with an AbTag cocktail targeting 154 unique cell surface antigens, including principal lineage markers, plus 9 isotype controls. (G) FICTURE projection of spatial factors identified via LDA analysis of protein expression results (G, upper). Twelve factors are annotated and color-coded (G, lower). White and gray boxed areas are further analyzed in (H) and (J), respectively. (H) Magnified view of the white boxed region from (G), showing spatial factor projections. (I) Spatial heat maps for selected individual factors in (H) demonstrate the positional probability of each factor. For spatial heat maps of all 12 LDA factors for a wider area, refer to Fig. S5E. (J) Magnified view of the gray boxed region from (G), showing additional spatial factor details. (K) A simplified diagram depicts the spatial arrangement of various B cell and T cell areas, such as marginal zone, T cell zone, and primary and secondary follicular areas, using the same color coding as in (G, H, and J). All scale bars indicate the scale adjusted to account for tissue expansion.

Fluorescence imaging of the expanded gel revealed that oligo-dT probes—which hybridize with poly(dA) and thus highlight ADT locations—were mostly localized around the cell membranes surrounding DAPI-stained nuclei (Fig. 5B). Notably, although the nuclear membrane was clearly visible in fluorescence imaging (Fig. 5B, right panel), it was less distinct in the Seq-Scope-X data (Fig. 5C, right panel). This discrepancy likely results from physical differences between the fluorescence imaging plane (in the middle of the gel) and the ADT capture interface (at the gel–array contact). Subtle subcellular details may have been obscured by diffusion of ADT molecules and their subsequent capture by spatial barcode molecules on the Seq-Scope Chip.

Nevertheless, mapping individual ADTs revealed highly localized expression patterns (Fig. 5D, S5A), indicating that Seq-Scope-X successfully captured spatial proteomic information with intact positional fidelity and a broad dynamic range. Using Latent Dirichlet Allocation (LDA)-based FICTURE analysis, we identified 12 distinct spatial factors representing diverse immune cell types and states (Fig. 5E, S5B). These factors included multiple T-and B-lymphocyte populations and myeloid-lineage cells (Fig. 5F, S5C). T and B cell zones were clearly separated, and within the T cell areas, CD4⁺ and CD8⁺ T cells were distinguishable, mirroring their tight intermingling observed in multispectral imaging [39]. While single-cell CITE-seq (scCITE-seq) [33, 34, 38] achieves similar molecular resolution in distinguishing these populations, it lacks spatial context. Conversely, current spatial proteomics platforms such as SM-Omics [35], SPOTS [36] and spatial CITE-seq [37] provide coarse spatial information but lack the resolution to reliably distinguish closely associated cell populations at the tissue level. We also observed rich diversity in the B cell compartment, with distinct cell populations expressing IgM, CD20, CD21, and CD79b, as well as antibody staining patterns indicating myeloid cells such as dendritic cells and macrophages (Fig. S5C).

These results establish that Seq-Scope-X exhibits both the single cell-level molecular precision of scCITE-seq and substantially higher spatial resolution than existing spatial proteomics technologies. While the current study utilized commercially available cell-surface antibody panels designed for single-cell CITE-seq, future applications could incorporate intracellular, extracellular matrix, or even post-translationally modified protein panels, enabling unlimited multiplexing capabilities in a single-step experiment.

### Seq-Scope-X Maps Human Tonsil Spatial Proteome

We next evaluated Seq-Scope-X’s ability to profile the cell surface proteome in human tonsil tissues undergoing chronic tonsillitis by using an antibody cocktail targeting 154 human cell surface antigens. Similar to the mouse spleen analysis, Seq-Scope-X identified a broad spectrum of T-and B-lymphocyte populations, as well as myeloid, epithelial, and endothelial cells (Fig. 5G, S5D–S5F). Notably, distinct B-cell populations were observed in primary and secondary follicle regions, as well as in extrafollicular areas (Fig. 5H, 5I). This pattern, which was consistent across multiple tissue regions (Fig. 5J, 5K), demonstrated the Seq-Scope-X’s capacity to resolve the complex organization of tonsillar secondary lymphoid tissue structures. The secondary follicle contained its characteristic mixture of dendritic cells and neutrophils, along with activated plasma cells expressing immunoglobulin light chain genes (Fig. 5I, S5F). Seq-Scope-X also accurately identified both CD8+ and CD4+ T cells which were largely present in the T cell areas adjacent to B follicles (Fig. S5E). Portions of the tissue additionally expressed epithelial and endothelial markers, indicating stromal regions adjacent to the lymphoid organ (Fig. S5E). These data confirm that Seq-Scope-X can reliably detect and spatially resolve proteomes in diverse tissues.

### Seq-Scope-X with DMAA Chemistry Enables Greater Expansion

All Seq-Scope-X data in Figures 1–5 was generated with standard expansion gels composed of acrylamide and acrylate, which could allow for up to fourfold tissue expansion in conditions allowing for RNA capture [24]. This translates to a maximum effective resolution of 150–300 nm (Fig. 6A). To push the resolution further, we incorporated X10 microscopy, which replaces acrylamide with dimethylacrylamide (DMAA) to achieve up to tenfold expansion [40–42]. In the X10 microscopy, DMAA’s unique polymerization properties produce a softer, more swellable hydrogel, enabling a maximum effective resolution of 60–180 nm (Fig. 6A), which is considered as a true nanoscale regime.

**Fig 6.**
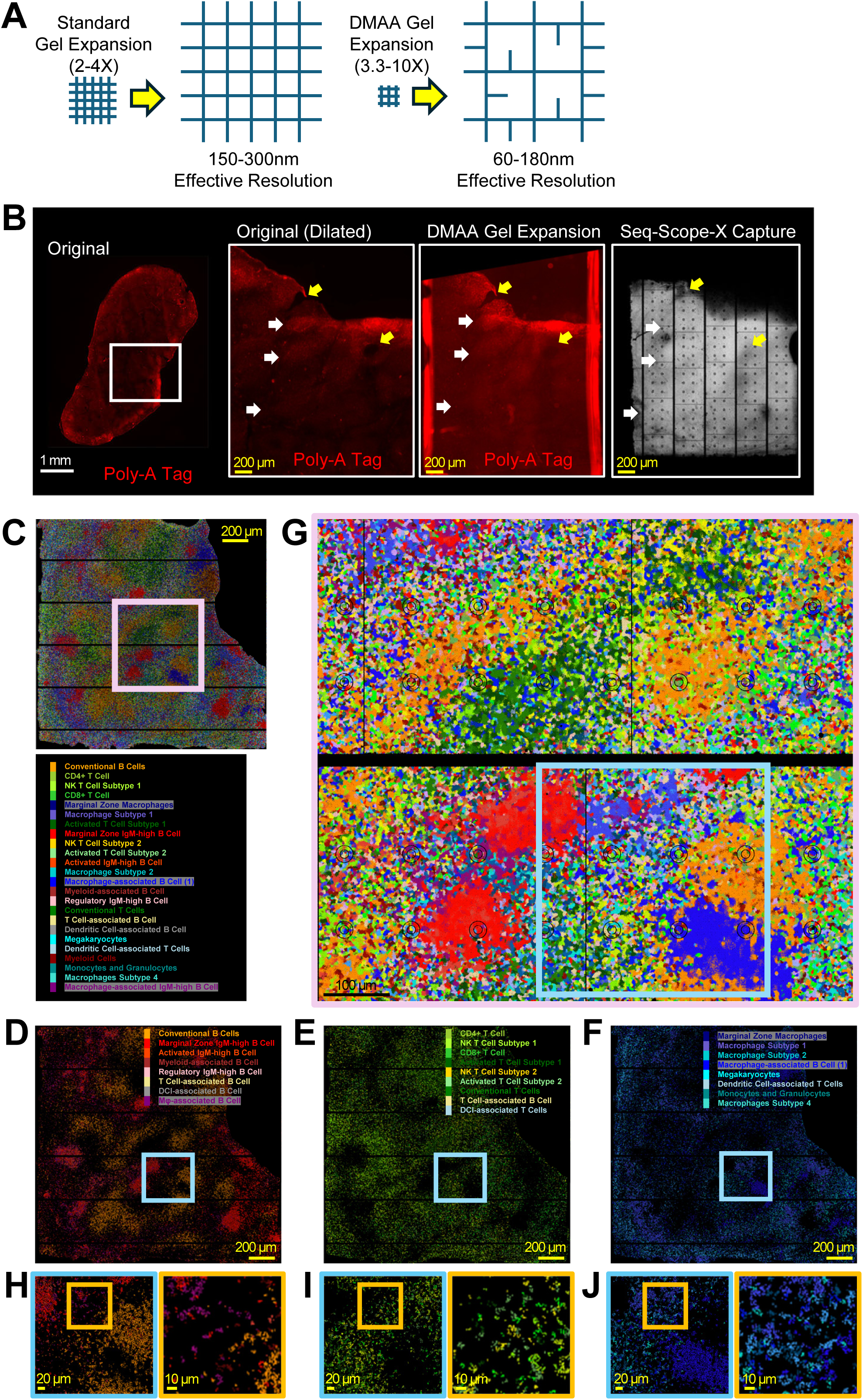
Seq-Scope-X is Compatible with DMAA Chemistry for Higher Resolution Analysis. (A) Schematic diagram comparing expansion chemistry using standard (left) and DMAA (right) gels. While not a precise representation of the final polymer arrangement, this schematic provides a conceptual visual guide to illustrate the differences between the two gels. The standard gel uses a separate backbone (acrylamide-acrylate) and crosslinker (methylenebisacrylamide), forming a more defined net structure. In contrast, the X10 gel employs dimethylacrylamide, which can function as both a backbone (with acrylate) and a crosslinker, resulting in a more flexible structure capable of gross expansion. (B-J) Seq-Scope-X analysis of the spatial proteome in mouse spleen, enhanced by DMAA gel expansion, using an antibody tag cocktail targeting 119 unique cell surface antigens, including principal lineage markers, plus 9 isotype controls. (B) Comparison of the spleen tissue used in the analysis, before and after tissue expansion. The spleen section was imaged for oligo-dT-acrydite fluorescence, which reflects the presence of Poly-A tags in the tissue. The boxed area is digitally magnified (left inset), and compared with the expanded image (center inset) and the digitally reconstructed image from Seq-Scope-X dataset (right inset). The dimensions and geometry of salient features (white and yellow arrows) are concordant across representations. (C-J) FICTURE projection of spatial factors identified via LDA analysis of protein expression results. Twenty-four factors are annotated and color-coded as shown in (C). The pink box in (C) is magnified in (G). Blue boxes in (D-G) are further magnified in (H-J, left). Yellow boxes in (H-J, left) are subsequently magnified in (H-J, right). Different B cell (D), T cell (E), and myeloid cell (F) subtypes are represented with reddish, greenish, and bluish colors, respectively. White scale bars represent the original scale, while yellow and black scale bars indicate the scale adjusted to account for tissue expansion.

To assess the compatibility of Seq-Scope-X with the DMAA expansion chemistry, we repeated the mouse spleen analysis (Fig. 5A–5F) using DMAA instead of acrylamide. This resulted in an approximately 3.3-fold expansion while preserving the salient spatial features of the original tissue (Fig. 6B, first inset; indicated in arrows), as validated in both the expanded gel (Fig. 6B, second inset) and the Seq-Scope-X array’s digital output (Fig. 6B, third inset). FICTURE analysis of the Seq-Scope-X dataset produced from the DMAA gel recapitulated all major cell types and histological regions identified in the standard Seq-Scope-X procedure, while providing enhanced spatial detail (Fig. 6C). In particular, it revealed a wider diversity of B-cell (Fig. 6D), T-cell (Fig. 6E), and myeloid cell (Fig. 6F) subtypes, along with their specific marker expression (Fig. S6A) and spatial distributions (Fig. S6B). Furthermore, digital magnification of the spatial factor map demonstrated clear, cellular-level organization of these immune populations (Fig. 6G–6J), confirming that Seq-Scope-X could be successfully combined with the DMAA expansion chemistry, which could open a new way of further increasing the resolution to enable nanoscale spatial analysis.

## Discussion

The development of Seq-Scope-X represents a significant advancement in sequencing-based spatial omics, by closing the resolution gap between sST and iST methodologies. By integrating tissue expansion with Seq-Scope, Seq-Scope-X enhances spatial precision, transcriptomic coverage, and compatibility with multiplexed proteomic analyses, enabling detailed subcellular and cellular investigations across various tissues.

In this study, we demonstrated that Seq-Scope-X could achieve a resolution up to ∼180 nm—higher than most conventional light microscopes. In terms of pixel density, this translates to an order-of-magnitude improvement over standard Seq-Scope (from ∼3 million pixels per mm² to ∼30 million pixels per mm²). When tissue expansion is further optimized and maximized, resolutions of ∼60 nm is theoretically possible with the DMAA expansion chemistry, further improving upon what we have already achieved. Even with a modest 3–3.3× tissue expansion demonstrated in the current work, Seq-Scope-X enables precise mapping of subcellular transcriptomic heterogeneity. The ability to resolve such features brings Seq-Scope-X’s resolution closer to that of iST technologies like Xenium, MERFISH and CosMx, while maintaining the key advantages of sequencing-based approaches: comprehensive whole-transcriptome coverage and superior scalability in both time and cost. Additionally, because mRNA transcripts are captured and immobilized prior to tissue expansion, the effective lateral diffusion is diminished in Seq-Scope-X relative to standard Seq-Scope, scaling inversely with the expansion factor. Indeed, our analyses of liver, brain, and colon tissues confirm Seq-Scope-X’s superior spatial resolution and precision, which are critical for understanding subcellular structures, organelle distributions, zonation patterns, and other fine-scale cellular and tissue-level properties.

The power of Seq-Scope-X to reveal previously inaccessible biological insights is exemplified by its analysis of the liver transcriptome. While recent single-cell and spatial studies have established that hepatocytes express distinct gene sets based on their histological niches [25, 32], effectively dividing the liver’s metabolic labor, the flexibility of these metabolic roles remained unknown. Through systematic analysis of the Seq-Scope-X dataset, using both segmentation-based nuclear/cytoplasmic analyses and segmentation-free FICTURE approaches, we uncovered widespread mismatches between nuclear and cytoplasmic transcriptomic profiles within individual cells. These observations were independently validated through orthogonal imaging-based methods, including MERFISH and smFISH. Our findings suggest that hepatocellular characteristics are not fixed, but rather hepatocytes can dynamically switch their metabolic roles over time.

Seq-Scope-X’s capacity to separate nuclear and cytoplasmic transcriptomes offers new opportunities to improve cell trajectory analyses. Traditional approaches frequently rely on splicing states, where spliced transcripts represent a cell’s immediate function and unspliced transcripts suggest preparatory regulatory directions [43, 44]. However, splicing-based methods can be confounded by substantial gene-specific differences in splicing dynamics, raising concerns about their reliability in depicting temporal changes [44–46]. In contrast, Seq-Scope-X delineates transcriptomic properties into nuclear (future) and cytoplasmic (current) compartments, sidestepping the biases introduced by splicing kinetics. Our observations show that while unspliced and spliced reads from the same cell often diverge in clustering space (Fig. S2F-S2I), nuclear and cytoplasmic reads from the same cell integrate well (Fig. 2E), differentiating only in transcripts known to be nuclear-or mitochondria-specific (e.g., *Malat1*, *Neat1* and mitochondria-encoded genes).

This nuclear-cytoplasmic segmentation is most effective in larger, regularly shaped cells like hepatocytes, which have clear nuclear and cytoplasmic boundaries. Smaller or irregular cells (e.g., immune cells, cholangiocytes) pose additional challenges, as do inherently complex tissues such as the brain and colon, where precise subcellular segmentation remains more difficult—even though nuclear-specific factors can still be detected by unbiased clustering. Furthermore, our current liver data reflect a stable metabolic state in conventionally reared mice; the semi-random mismatches we observed between nuclear and cytoplasmic phenotypes may represent fine-grained fluctuations rather than broader shifts driven by metabolic cues. In future studies, feeding, fasting, hormone treatments (e.g., insulin or glucagon), hepatotoxic exposures, or partial hepatectomy could be used to reveal dynamic changes in nuclear vs. cytoplasmic transcriptomes and to track directional processes in liver zonation or tissue regeneration. Also, further optimization of the DMAA expansion chemistry within the Seq-Scope-X workflow could enable higher nanoscale (<100 nm) resolution, potentially extending nuclear-cytoplasmic segmentation to non-hepatic tissues and resolving even smaller subcellular features such as RNA granules.

Beyond transcriptomics, Seq-Scope-X can detect any poly-A–labeled molecule, allowing us to demonstrate single-cell–level proteomic analyses via nucleotide-tagged antibodies. Although we employed a cocktail designed for single-cell CITE-seq (thus targeting cell surface proteins), Seq-Scope-X can, in principle, detect intracellular and extracellular matrix proteins as well. A case in point is IgM: in plasma cells, it typically resides in the ER or is secreted into surrounding tissue. Because single-cell CITE-seq focuses on surface markers, IgM-rich plasma cells were not identified in prior datasets, whereas Seq-Scope-X detected them by capturing intracellular IgM. The approach may also capture secreted IgM within the local microenvironment. As more antibodies for intracellular or matrix-associated targets become available, a broader range of proteins and their post-translational modifications could be visualized at nanoscale resolution. Moreover, this approach presents a new tool to identify novel cell populations with unconventional marker protein expression and potentially at unexpected locations. When combined with single cell-level sequencing, the transcriptomic and genomic features of these novel cell populations, as well as their histological niche in the microenvironment, can be additionally revealed by this approach.

Seq-Scope-X’s strategy of separating tissue staining from spatial array capture confers several advantages over existing array-based spatial proteomics platforms (e.g., SM-OMICs [35] and SPOTs [36]), which rely on low-resolution arrays and apply antibodies directly to the capture surface. This can result in non-specific capture of antibody barcodes leading to undesirable noises. While microfluidic-based techniques, such as spatial CITE-seq/DBiT-Seq [37], or slide maneuvering-based techniques, such as 10x Visium CytAssist [20], mitigate nonspecific capture, they still operate at lower resolution (>2–20 µm). In contrast, Seq-Scope-X capitalizes on the high resolution of Seq-Scope augmented by tissue expansion, achieving nanoscale omics exploration with greater precision.

Separating tissue preparation from spatial capture further broadens Seq-Scope-X’s versatility. Various poly-A–labeled probes—such as wheat germ agglutinin, phalloidin, aptamers, or RNA in situ probes— can be applied without risking damage to the capture array. It is even conceivable to integrate genomic assays (e.g., as demonstrated in spatial genomics [47], spatial-ATAC [48, 49] and spatial-Cut&Tag [50, 51]) by labeling fragmented DNA with poly-A or other capture sequences, thereby extending high-resolution mapping to epigenomic states and transcription factor binding sites on a solid-phase platform.

Despite its groundbreaking capabilities, Seq-Scope-X has limitations requiring further refinement. Background signals from nonspecific antibody binding remain a challenge, common to most immunostaining workflows. Proximity labeling strategies, as employed in both O-Link bulk proteomics [52] and Visium FFPE [20], could mitigate this issue by requiring two probes to bind in close proximity to a target before generating a detectable signal. This ensures that only specific interactions contribute to spatial data, minimizing noise. Similarly, the reliance on temperature-based melting for poly-A release imposes potential problems with efficiency and diffusion artifacts; alternatives like photocleavable linkers or non–poly-A sequences may offer greater flexibility. Additionally, while the use of DMAA expansion chemistry enhances resolution, the theoretical tenfold expansion has not yet been fully achieved, leaving room for optimization. Current capture efficiency is comparable to standard Seq-Scope but requires improvement for deeper, more reliable transcriptomic analyses, especially in small or complex cell types where spatial footprints are limited.

In conclusion, Seq-Scope-X provides a transformative leap in spatial omics, attaining nanoscale resolution and enabling integrative transcriptomic and proteomic analyses at subcellular precision. Its adaptability, high accuracy, and multi-omic potential make it a valuable platform for probing tissue organization, cellular regulation, and disease pathogenesis. As future enhancements address the remaining technical challenges, Seq-Scope-X has the potential to establish itself as a foundational tool in high-resolution spatial biology.

## EXPERIMENTAL PROCEDURES

### Tissue Samples

Mouse tissues, including liver, colon, and spleen, were collected from 8-week-old male C57BL/6 wild-type mice, and brain tissue was obtained from a 6-month-old male mouse. All mouse experiments were conducted in accordance with protocols approved by the Institutional Animal Care and Use Committee (IACUC) at the University of Michigan.

Human tonsil tissue was obtained from the UCLA Pathology Biobank as a discarded surgical specimen from an adult patient diagnosed with chronic tonsillitis. The sample was fully de-identified prior to acquisition and is classified as IRB-exempt under institutional and federal guidelines. No identifiable patient information was accessible, and no additional ethical review or consent was required.

### Tissue Sectioning, Attachment, Fixation and Permeabilization

Tissue expansion techniques, previously used to enhance the resolution of iST [21, 53, 54] and low-resolution sST [23] approaches, were adapted for Seq-Scope-X analysis. Fresh frozen tissues embedded in Optimal Cutting Temperature compound (OCT) were sectioned using a cryostat (Epredia, 957250L CryoStar NX50) at-17°C, with a 5° cutting angle and a thickness of 10–16 µm. The tissue sections were mounted on charged microscope slides and fixed for 10 minutes at room temperature in 4% formaldehyde solution prepared by diluting 16% formaldehyde (28908, Thermo Scientific) in 1X PBS (14190, GIBCO). After fixation, tissue sections were washed three times with 1X PBS to remove residual formaldehyde.

The tissues were then permeabilized with ice-cold methanol at-20°C for 20–30 minutes, followed by three washes in 2X saline sodium citrate (SSC), prepared by diluting 20X SSC stock solution (BP1325, Fisher Bioreagents) with UltraPure water (10977, Invitrogen).

### Probe Hybridization

Tissues were incubated at room temperature in 30% formamide (AM9342, Invitrogen) diluted in 2X SSC for 30 minutes, then blow-dried using nitrogen gas. The anchor probe hybridization solution was prepared with 2X SSC, 30% formamide, 1 mg/mL Yeast tRNA (15401011, Invitrogen), 10% (w/v) dextran sulfate (J63606, Thermo Scientific Chemicals), 1% (v/v) SUPERase-In (AM2696, Invitrogen), and 2 mM Anchor-590ST (/5Acryd/TT+T T+TT +TT+T T+TT +TT+T /3ATTO590N/ as an HPLC-purified oligonucleotide; Integrated DNA Technologies) in UltraPure water (10977, Invitrogen). Dried tissues were submerged in the hybridization solution, covered with an RNase Zap (AM9780, Invitrogen)-treated coverslip, and incubated in a pre-warmed humidification chamber at 37°C for 48 hours.

### Tissue Embedding

The monomer solution was prepared as 2 M NaCl, 12% (w/v) sodium acrylate (408220, Sigma-Aldrich), 3% (v/v) 19:1 acrylamide/bis-acrylamide solution (BP1406, Fisher Chemical), and 60 mM Tris-HCl (15568, Invitrogen) in UltraPure water, followed by sonication at 40 kHz for 10–20 minutes (Branson, model no.1800). After disassembling humidified chamber, hybridized samples were washed three times with 2X SSC and then incubated in the monomer solution containing 0.2% tetramethylethylenediamine (TEMED, 17919, Thermo Scientific) at 4°C for 30–45 minutes. After removing the initial monomer solution, a polymerization chamber was constructed using a spacer (SLF-0601, Bio-Rad) with open ends for solution entry and a Gel Slick (50640, Lonza)-treated coverslip. The monomer solution was aliquoted and polymerization was initiated by adding 2% ammonium persulfate (0486, VWR) and 0.2% TEMED. The pre-polymerized solution was applied to the tissue, with capillary pressure between the glass slide and coverslip maintaining the solution in place. Polymerization was completed at room temperature within 1–2 hours.

### Tissue Clearing

The tissue digestion buffer was prepared as 2X SSC, 2% (w/v) sodium dodecyl sulfate (428015, Millipore), and 0.5% Triton-X (M143, AMRESCO) in UltraPure water. After polymerization, the slide-coverslip was disassembled to obtain the gel, which was trimmed to the desired size. The gel, still attached to the coverslip, was placed in a petri dish and washed three times with the digestion buffer.

Then the gel was incubated with the digestion buffer containing 1% Proteinase K (P8107S, NEB) to facilitate tissue clearing. The petri dish was then placed in a 37°C humidity chamber for 24 hours.

### Tissue Expansion

The digestion buffer was removed from the petri dish, and the gel was washed with 1X SSC. Tissue expansion was performed by incubating the gels in 0.1X SSC. The solution was refreshed after 15 minutes and replaced again after another 15 minutes. Following this, the gels were incubated for an additional ∼30 minutes or until the desired scale factors were achieved, depending on the tissue expansion rate. During the final 30 minutes of incubation, DAPI (D21490, Invitrogen) was added to assess tissue integrity and assist with tissue alignment.

### Imaging & Probe Release

Expanded tissue gels were trimmed with a razor to fit the ∼7 mm x 7 mm dimensions of the Seq-Scope Chip, produced as described in the previous Seq-Scope protocol [15]. DAPI and poly-dT probe (ATTO590) images of the expanded tissue samples were captured in a digital darkroom (Keyence BZ-X810). To minimize gel drying, distortion, or RNA damage, imaging was often skipped or performed at low magnification (e.g., using a 4X objective lens). When captured, the DAPI and poly-dT images were used for downstream alignment and segmentation. Following imaging, the gels were incubated in a RapidFISH slide hybridizer (240200, Boekel Scientific™) at 45°C for 30 minutes. The heat facilitated the release of RNAs or antibody tags from the anchor probes, which were subsequently captured by the HDMI-oligo-dT probes on the surface of the Seq-Scope Chip.

### Standard Procedure and Library Preparation

After incubation, the gel was carefully removed from the sandwich to retrieve the Seq-Scope Chip with spatially captured RNAs. The established Seq-Scope protocol [15] was then followed, starting from the reverse transcription step. All subsequent steps described in the Seq-Scope protocol [15] were strictly adhered to, except for the tissue digestion step (Proteinase K treatment), which was omitted as no real tissue was attached to the Seq-Scope Chip.

### Tissue Processing and Antibody Staining for Seq-Scope-X Proteome Analysis

For Seq-Scope-X proteome analysis, tissues were permeabilized with ice-cold methanol at-20°C for 20 minutes. Following permeabilization, tissues were washed with PBS containing 1% Triton-X (Ab-wash solution) and incubated with a blocking solution consisting of 1% BSA (BP1605, Fisher Scientific) and 1% RNase inhibitor (30281, Lucigen) in Ab-wash solution for 1 hour at room temperature. The TotalSeq™-A Human (399907, BioLegend) and Mouse (199901, BioLegend) Universal Cocktail were reconstituted according to the manufacturer’s recommendation and diluted in SignalStain® Antibody Diluent (8112, Cell Signaling). The tissues were incubated with the diluted antibody cocktail overnight at 4°C. After incubation, the antibody solution was removed, and the tissues were washed three times with the Ab-wash solution for 5 minutes each. Finally, the tissues were subjected to post-fixation with 4% formaldehyde in PBS for 10 minutes to stabilize the staining.

### Tissue Expansion and Library Preparation for Seq-Scope-X Proteome Analysis

After post-fixation, the antibody-stained slides were processed following the same steps as the Seq-Scope-X transcriptome analysis, including “Probe Hybridization,” “Tissue Embedding,” “Tissue Clearing,” “Tissue Expansion,” and “Imaging & Probe Release.” However, for Seq-Scope-X proteome analysis, specific modifications were made to capture the DNA oligonucleotides tagging TotalSeq™-A antibodies. The antibody-tagging DNA oligonucleotides encode small RNA Read 2 (SR2) PCR adapter, antibody tag (AT) barcode, a random nucleotide (used as unique molecular identifier (UMI) in our analysis) and poly-dA sequences. Following the “Imaging & Probe Release” step, reverse transcription was performed as in Seq-Scope-X transcriptome analysis. During this step, reverse transcriptase, with its DNA polymerase activity, extended the antibody-tagging DNA oligonucleotides from its poly-dA tail using the Seq-Scope Chip’s DNA template encoding oligo-dT, HDMI and Truseq Read 1 (TR1) adapter sequences. Consequently, the primary antibody tag (AT) library, containing SR2, Ab-barcode, UMI, poly-A, HDMI and TR1 sequences, were produced on the Seq-Scope Chip. The reverse transcription solution was removed from the Chip, and the primary AT library was eluted using 30 μL of 0.1 N NaOH for 5 minutes. This elution step was repeated twice to collect a total of 60 μL of elution solution, which was neutralized with 30 μL of 3 M potassium acetate, pH 5.5 (AM9610, Invitrogen). The neutralized primary AT library was purified using the Oligo Clean & Concentrator kit (D4060, Zymo Research) per the manufacturer’s instructions, yielding a 40 μL final elution volume. The primary AT library was then amplified using an AT-PCR reaction comprising 100 μL of 2X Kapa HiFi Hotstart ReadyMix (KK2602, KAPA Biosystems), 1 μL each of 100 mM RPEPCR*F (TCT TTC CCT ACA CGA CGC*T*C) and SR2-RPEPCR*R (CCT TGG CAC CCG AGA ATT C*C*A) primers, 40 μL of the purified primary library, and 58 μL of Ultrapure water. The AT-PCR cycle condition is: 95°C for 3 minutes, followed by 15 cycles of 95°C for 30 seconds, 60°C for 1 minute, 72°C for 1 minute, and a final extension at 72°C for 2 minutes. The AT-PCR product was purified using AMPure XP beads (A63881, Beckman Coulter) with a 1.8X bead/sample ratio and eluted in 30 μL. The sequencing library was prepared in an AT-Indexing-PCR reaction using 20 μL of 2X Kapa HiFi Hotstart ReadyMix, 4 μL each of 10 mM WTA1*F (AAT GAT ACG GCG ACC ACC GAG ATC TAC AC [10-mer index sequence] TCT TTC CCT ACA CGA CGC TCT*T *C) and SR2-WTA1*R (CAA GCA GAA GAC GGC ATA CGA GAT [10-mer index sequence] GTG ACT GGA GTT CCT TGG CAC CCG AGA ATT C*C*A) primers, 3 μL of the purified AT-PCR product, and 9 μL of Ultrapure water. The AT-Indexing-PCR cycle condition is: 95°C for 3 minutes, followed by 6–8 cycles of 95°C for 30 seconds (8 cycles for >0.2 ng/μL, 7 cycles for >0.5 ng/μL, and 6 cycles for >0.8 ng/μL), 60°C for 1 minute, 72°C for 1 minute, and a final extension at 72°C for 2 minutes. The AT-Indexing-PCR products were purified using AMPure XP beads with a 1X bead/sample ratio and eluted in 30 μL. The purified AT-Indexing-PCR products underwent size selection via agarose gel elution to isolate the 250 bp band using the Zymoclean Gel DNA Recovery Kit (D4001, Zymo Research) according to the manufacturer’s instructions. The final library was sequenced using an Illumina NovaSeq-X sequencer.

### Tissue Expansion with DMAA Gels

Tissue expansion using the X10 system was achieved with Dimethyl Acrylamide (DMAA) gels, as previously described for expansion microscopy [41]. In this method, the Acryl/bis-acrylamide used in the standard monomer solution was replaced with DMAA (274135, Sigma Aldrich) to allow for larger expansion. The DMAA monomer solution was prepared following the X10 expansion microscopy protocol [41], with modifications to include 80% DMAA, 20% sodium acrylate, 2 M NaCl, and 60 mM Tris-HCl (pH 8.0). These adjustments optimized expansion speed and stabilized molecular bonding between the anchor probe and mRNA/antibody tags. The expansion factor of DMAA gels was dependent on the volume of 0.1X SSC used during the process. Larger volumes increased the expansion factor (up to 10×, as described in the original protocol [41]), but excessive expansion complicated gel handling. To achieve a balance, a volume between 5 mL and 10 mL was chosen, resulting in a modest expansion factor of approximately 3.3×.

## QUANTIFICATION AND STATISTICAL ANALYSIS

### Seq-Scope-X Transcriptome Data Processing

Seq-Scope-X data analysis was performed using the same pipeline as the Seq-Scope pipeline [15] with standard parameters and adjustments for the experimentally determined scale factor. The analysis followed the steps outlined in the NovaScope and NovaScope Exemplary Downstream Analysis (NEDA) documentation, which include the generation of spatial barcode maps, alignment of sequencing reads to reference genomes, creation of spatially resolved gene expression (SGE) matrices, hexagonal binning, Seurat-based clustering [55], Latent Dirichlet Allocation (LDA)-based spatial factor discovery, pixel-level spatial factor projection using FICTURE [30], alignment of transcriptome data with microscopic imaging data, and image-based segmentation analyses [12]. The fluorescence images, such as DAPI and oligo-dT fluorophore images, did not have fiducial marks, so we aligned them to the Seq-Scope-X transcriptome image with salient features, such as blood vessel locations and nuclear centers estimated by unspliced RNAs, using the georeferencing function of QGIS. The aligned image was further enhanced using CellProfiler and Photoshop to clearly indicate the nuclear or RNA/ADT-captured area without the background signal. The Seurat v5 package [55] was used to analyze the hexagonally binned data through PCA and clustering analyses, as well as differential expression analyses. FICTURE was also used to project seurat clusters into Seq-Scope raw pixel space, in the NEDA workflow. Additional analyses not performed on this pipeline and packages are outlined separately below.

### Resolution and Pixel Density Benchmarking

For resolution benchmarking in Fig. 1E, Seq-Scope-X was compared with 11 prior technologies. The center-to-center resolution of each method was as follows: ST (200 μm) [6], 10x Visium (100 μm) [7], Slide-Seq (10 μm) [8, 9], HDST (4 μm) [11], DBiT-Seq (20 μm) [27], Seq-Scope (0.5 μm) [12], Stereo-Seq (0.5 μm) [13], Pixel-Seq (1 μm) [14], Expansion ST (40 μm) [23], 10x Visium HD (2 μm) [20], and Open-ST (0.6 μm; this resolution point also represents Seq-Scope^NOVASEQ^ and Nova-ST) [15–17]. These values were obtained as nominal resolutions from their respective publications. Seq-Scope-X demonstrated a resolution of 0.2 μm, achieved through a 3× expansion factor, as validated using the mouse liver dataset. Even though it is not included in Fig. 1E, incorporation of the DMAA chemistry into Seq-Scope-X achieved 3.3× expansion factor and 180 nm resolution and can potentially achieve up to a 6-10× expansion factor and 60-100 nm. These reflect an expansion-mediated enhancement from the 0.6 μm resolution of NovaSeq-based Seq-Scope. Please note that the feature size of 10x Visium and Stereo-Seq is 55 μm and 0.2 μm; however, to perform the fair comparison between technologies, we only considered center-to-center distances for this benchmarking comparison.

For pixel density benchmarking in Fig. 1F, the following values in barcode/mm² were used: ST (25), Visium (154), Slide-Seq (2,210), HDST (94,100), DBiT-Seq (2,500), Seq-Scope (1,000,000), Stereo-Seq (4,000,000), Pixel-Seq (1,000,000), Expansion ST (625), Visium HD (250,000), Open-ST (2,961,000), and Seq-Scope-X (26,649,000). The method for calculating pixel density is done corresponding to our previous study [12]. In brief, for the technologies that have a defined pixel area (ST, Visium, HDST, DBiT-Seq, Stereo-Seq, Expansion ST, Visium HD, Open-ST, and Seq-Scope-X), pixel density was calculated as the inverse of the pixel area, adjusted for the expansion factor (Expansion ST & Seq-Scope-X). Note that HDST, Stereo-Seq, Open-ST, and Seq-Scope-X technologies employed patterned flow cells with a defined resolution and pixel density; however, due to sequencing errors, effective pixel density could be lower than the nominal density. For Slide-Seq, Slide-SeqV2 and Seq-Scope, pixel density was calculated in 150 μm grids (Slide-Seq and Slide-SeqV2) and 10 μm grids (original Seq-Scope) of the final dataset. Pixel-Seq pixel density was estimated to be the same as the original Seq-Scope as they are based on the similar sequencing chemistry.

Also note that Open-ST, Nova-ST, and NovaSeq-version of Seq-Scope were all published in 2024 with largely similar methodologies with minor differences in experimental and data processing procedures [15–17]. Their performances are similar to each other; so for simplicity, we only indicated Open-ST, which first appeared in the literature, in the graph.

For technologies developed from academic space, the date corresponds to either their peer-reviewed publication date or the preprint release date if not yet published. For commercial technologies, the date reflects when the technology became publicly and commercially available.

### RNA Capture Benchmarking and Transcriptome Comparison

To objectively and fairly benchmark transcriptome capture efficiency and precision, we compared Seq-Scope-X liver datasets with publicly available datasets derived from the same type of tissue, specifically normal young (2-4 months of age) wild-type male C57BL/6 mouse liver, ensuring consistency in biological context. These datasets included 10x Visium [26], xDBiT-Seq [27], Stereo-Seq [28], Seq-Scope^MISEQ^ [12], and Seq-Scope^NOVASEQ^ [15]. Additional datasets, including ST [25], which analyzed normal young (2-3 months of age) wild-type female C57BL/6 mouse liver, and Slide-Seq (v1) [8], which analyzed normal mouse liver of an unspecified origin, were also analyzed. For ST, Visium, Slide-Seq, and xDBiT-Seq, transcriptome capture efficiency per spatial feature (UMI/feature) was calculated and multiplied by spatial feature density (feature/μm²). For higher-resolution technologies such as Stereo-Seq and Seq-Scope, transcriptome capture efficiency was calculated using publicly available binned datasets (50 µm square grid for Stereo-Seq, 10 µm square grid for Seq-Scope^MISEQ^, and 14 µm hexagonal grid for Seq-Scope^NOVASEQ^) as UMI/bin divided by bin area (μm²/bin). Seq-Scope-X data was binned using a 5 µm-sided hexagonal grid (14 µm if not accounted for 3× expansion factor) and processed similarly. Outliers or non-tissue regions were removed using custom thresholds, such as minimum or maximum feature counts. Summary statistics for UMI counts were computed, and violin plots of UMI per unit area (UMI/μm²) were generated for comparison across datasets. To visualize spatial distribution and data quality, XY scatterplots stratified by raw (low-resolution ST) or binned (high-resolution ST) dataset were generated, with each spatial coordinate point colored by UMI count (nCount), allowing comparison of transcriptome capture across spatial features and datasets.

To assess transcriptomic similarity across datasets, we performed a comprehensive comparison using pseudo-bulk gene expression profiles aggregated from each ST dataset, along with genuine bulk RNA-seq results (six biological replicates) previously generated from the same type of tissue (normal wild-type C57BL/6 male mouse liver) [29]. Sum counts of gene expression from Seq-Scope-X and other datasets (ST, Visium, Slide-Seq, xDBiT, Stereo-Seq, Seq-Scope^MISEQ^, Seq-Scope^NOVASEQ^, and bulk RNA-seq samples) were merged based on shared genes. Transcriptome-wide similarity was visualized through scatterplots and heat maps of pairwise comparisons (Fig. S2A, S2B). These analyses underscore the consistency and reliability of Seq-Scope-X’s transcriptome data relative to prior methods.

Comparison of spatial gene expression, presented in Fig. 3G, 3H, 4B and 4C, were conducted by plotting each individual digital gene expression as colored dots. Following gene list, retrieved from clustering or factoring results as well as area-specific differential expression analyses, were used for spatial plotting. Fig. 3G and 3H: *Mup20* (blue), *Hamp* (green), *Mup17* (orange) and *Glul* (red). Fig. 4B: *C1ql3*, *Cntnap5a*, *Dpyd*, *Glis3*, *Htr4*, *Il1rap*, *Itga8*, *Jph1*, *Lrrtm4*, *Maml2*, *Ppfia2*, *Prkd1*, *Prox1*, *Rasal2*, *Rfx3*, *Sema5a*, *Stxbp6*, *Tafa2* and *Trpc6* (DG Granular Cells, red), *Atp2b1*, *Cacnb2*, *Epha6*, *Galntl6*, *Hs6st3*, *Kcnh7*, *Man1a*, *Nr3c2*, *Ppm1e*, *Ryr3*, *Slc24a3* and *Slc8a1* (CA1, green), *Cacna2d3*, *Cpne4*, *Kcnq5*, *Nrip3*, *Tafa1*, *Wipf3*, *Zfp804a*, *Cntnap5c*, *Hpca*, *Khdrbs3*, *Neto1*, *Sv2b*, *Tafa1* and *Ywhah* (CA2/3, yellow), *Chgb*, *Epha3*, *Lmo4*, *Neurod6*, *Nptxr*, *Nrp1*, *Rnf182*, *Slit2*, *Trhde* and *Trps1* (CA3, orange), *Adarb1*, *Amotl1*, *Kcnc2*, *Lhfp*, *Nefh*, *Nell1*, *Nexn*, *Ntng1*, *Pde7b*, *Pdp1*, *Prkcd*, *Rora*, *Shox2*, *Slc17a6*, *Synpo2*, *Tcf7l2* and *Zfhx3* (Excitatory, lightblue), *Adam18*, *Adam32*, *Aip*, *Akr1c14*, *Ankrd52*, *Arrdc3*, *Atad5*, *Bcar3*, *Bdh1*, *C2cd4c*, *Cacna1i*, *Cct3*, *Cdh18*, *Cntnap4*, *Col12a1*, *Commd6*, *Csrnp2*, *Ctso*, *Dcbld1*, *Ddt*, *Dhrs3*, *Dnaaf11*, *Dpp10*, *Dzip1l*, *ENSMUSG00000120124*, *Efhd1*, *Fam124b*, *Ggh*, *Gm10033*, *Gm13322*, *Gm5468*, *Gm57059*, *Gpn3*, *Gpr176*, *Hdac11*, *Hey2*, *Ipp*, *Josd1*, *Kcnj16*, *Kirrel*, *Lrrc61*, *Ltbp1*, *Lzts1*, *Maip1*, *Med12*, *Mtmr2*, *Nrg3os*, *Nxt2*, *Pdzrn4*, *Peli3*, *Phkg2*, *Pik3r3*, *Pms1*, *Ppargc1b*, *Ppil2*, *Ppp2r1b*, *Prdm15*, *Prox1os*, *Ptpn20*, *Rnd3*, *Rnf121*, *Sdad1*, *Sema5b*, *Septin9*, *Sh3yl1*, *Syt2*, *Tent4a*, *Tmem132a*, *Tmem196*, *Tmem268*, *Tox*, *Tspan31*, *Unk*, *Vwc2l*, *Zbtb25*, *Zfp68*, *Zfp703*, *Zfp715*, *Zfp763*, *Zfp982* and *Zxdb* (Astrocyte, violet), *Erbb4*, *Pcdh9*, *Pde4b*, *Plp1*, *Qki*, *Tmeff2* and *Tspan2* (Oligodendrocyte, blue), *Ttr* and *Enpp1* (Choroid Plexus, white). Fig. 4C: *Acta2*, *Actg2*, *Cnn1*, *Csrp1*, *Des*, *Flna*, *Hspb1*, *Myh11*, *Myl9*, *Mylk*, *Tagln*, *Tpm1* and *Tpm2* (Smooth Muscle, white), *Agr2*, *Cryab* and *Muc2* (Goblet I (DCSC), red), *Zg16* and *Fcgbp* (Goblet II, orange), *Mptx1* (Paneth, purple), *Igha*, *Jchain* and *Ly6e* (Immune, yellow), *S100a6* (Fibroblast, blue), *Saa1* and *Aqp8* (Colonocyte I, cyan), *Krt20*, *Cldn4* and *Fabp2* (Colonocyte II, green).

### Image-Based Single Cell and Subcellular Segmentation

Image-based segmentation for single cells was performed as described in NEDA documentation and our previous protocol [15], using the Watershed algorithm implemented in the NIH ImageJ and NEDA pipeline, with the following modifications. Instead of using the hematoxylin-eosin image that was exemplified in our previous protocol [15], we used the spliced (red) and unspliced (green) RNA density image, processed through Gaussian filtering (as in Fig. 1I and 2A), for image-based cell segmentation. When this image was converted to grayscale, green unspliced signal is more pronounced and suitable to identify local maxima, which is the nuclear center, while the dimmer red spliced signal will be still effectively used to estimate the cell boundary as spliced transcripts are scarcely found in extracellular areas. Each Watershed segments are then considered as a single cell, and a circular area centered at the local maxima with a radius of 5 µm (expansion-adjusted unit) is considered as a nuclear compartment, while the remainder of the single cell area was considered as a cytoplasmic compartment. Performance of this segmentation method is generally concordant with visual inspection of the image (as in Fig. 2A). For downstream analysis, each compartment was separately collapsed as independent aggregates, even though the data are labeled so that the aggregate pair from a single cell can be tracked for the cytoplasmic-nuclear mismatch analysis (as in Fig. 2G, 2H). The spliced and unspliced gene counts were retrieved using *velocyto* function in STARsolo, implemented in the NovaScope pipeline, and whole cell segment was used to aggregate single cell spliced and unspliced transcriptomes used for analysis in Fig. S2F-S2I.

### Analysis of In situ Imaging Data

MERFISH mouse liver data was retrieved from the Vizgen website [31] and subjected to FICTURE analysis as described previously [30] and analyzed with the background DAPI staining data after image alignment. Mouse smFISH liver data was retrieved from a previous publication [32], and reproduced with permission from Springer Nature [32]. These data demonstrate the discrepancies between nuclear and cytoplasmic hepatic gene expression related to metabolic zonation, an observation consistent with Seq-Scope-X findings.

### Seq-Scope-X Proteome Analysis

To enable proteome analysis, the NovaScope pipeline was modified to ingest the Read2 sequencing reads encoding 15 bp antibody tag (AT) barcode, a random nucleotide (used as UMI) and poly-dA sequences. Instead of running STARsolo for transcriptome alignment, a custom software (spatula match-tag, available at https://github.com/seqscope/spatula) was used to match the FASTQ reads with the antibody barcodes in the TotalSeq™-A Human Universal Cocktail, V1.0 (399907, Biolegend) and TotalSeq™-A Mouse Universal Cocktail, V1.0 (199901, Biolegend). The first 15 bases in Read 2 were used as antibody tags, and the next base was used as UMI. Sequence reads that do not match with any antibody barcodes are discarded, and the HDMI counts having the same AT barcode and UMI were deduplicated. The spatial digital gene expression (sDGE) output was constructed in a format similar to sDGE produced from transcriptome analysis, except that it only has one expression count and does not have intron-containing gene counts (GeneFull) or spliced/unspliced gene counts. The sDGE was used for downstream analysis in the same way as what was described in NEDA, such as LDA and FICTURE analyses [15].

## ACKNOWLEDGEMENTS

The authors thank the U-M Advanced Genomics Core (AGC) for their cooperation and help in performing Seq-Scope and sequencing analysis. We thank Lee, Han and Kang lab members for their help in experiments and analysis. The work was supported by the Taubman Institute (to H.M.K. and J.H.L.), NIH (T32AG000114 to A.A., C.S.C. and Y.S.K, UH3CA268091 and R01DK133448 to J.H.L., R01AG079163 to M.K. and J.H.L., and R01HG011031 to Y.S. and H.M.K.), Chan Zuckerberg Initiative (to H.M.K.), and Glenn Foundation (to J.H.L.) grants.

## DECLARATION OF INTERESTS

J.H.L is an inventor on a patent and pending patent applications related to Seq-Scope. H.M.K. owns stock in Regeneron Pharmaceuticals. Y.H. is currently an employee of Samsung Semiconductor. All other authors declare no competing interests.

---

**Fig S1.**
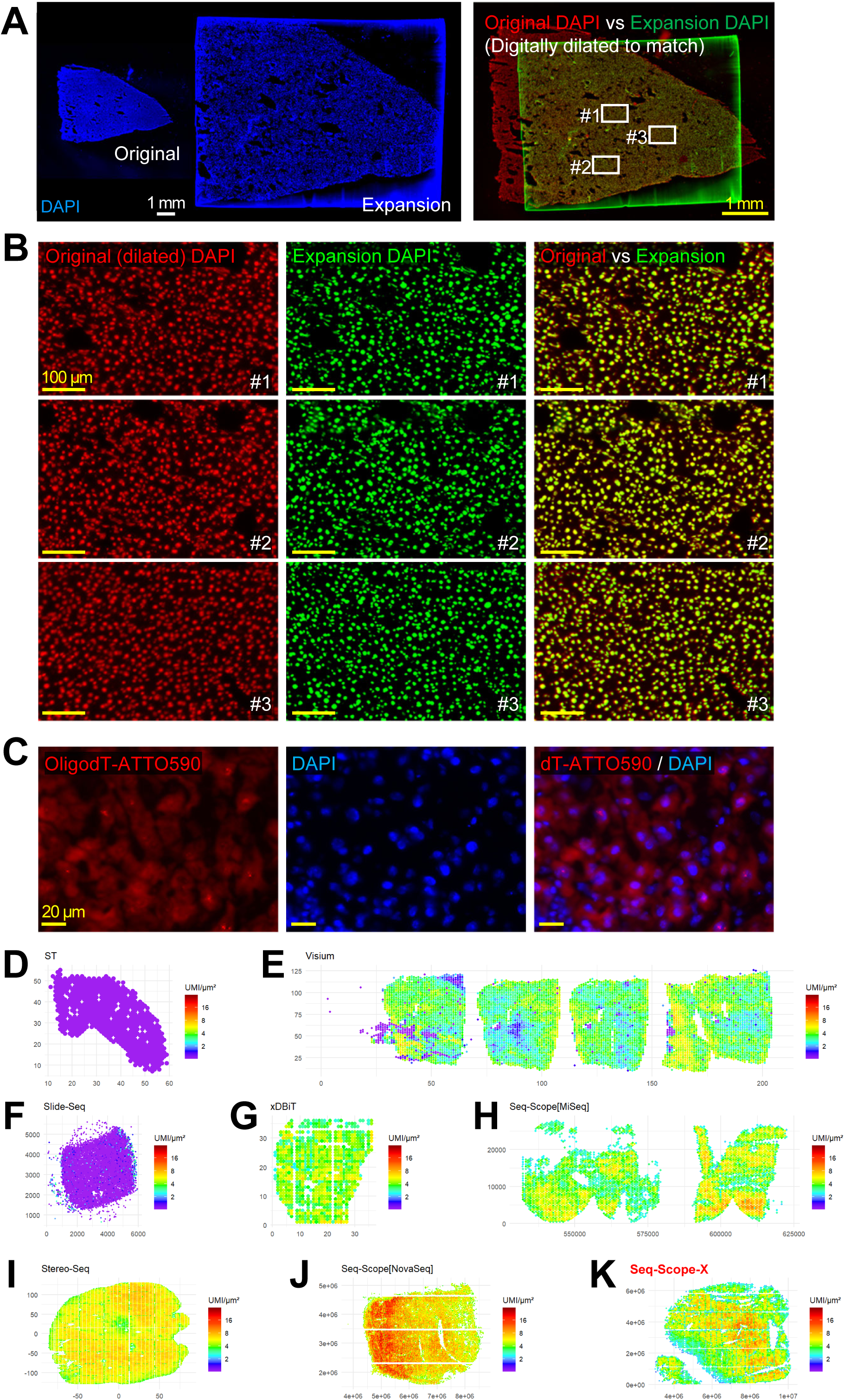
Tissue Expansion and Transcriptome Capture Performances in Seq-Scope-X. (A and B) Comparison of the liver tissue before and after tissue expansion. The liver section was imaged using conventional DAPI staining (Original). After hydrogel embedding and expansion (2.7X in the current data), the same tissue was imaged again (Expansion), and presented side-by-side to compare the scale (A, left). Each image was digitally colored into red (Original DAPI) and green (Expansion DAPI), and the original DAPI was digitally dilated to match the scale of expansion DAPI, and overlaid to each other. The overlaid image (A, right) demonstrates that the tissue expansion was uniformly applied to the whole tissue without any significant distortion or artifacts. Three regions (#1-#3 in A, right) were selected and further magnified (B) to show that microscopic structures were also well preserved before and after expansion. (C) Liver tissue was hybridized with ATTO590-labeled oligo-dT acrydite probe, embedded in an acrylite hydrogel, digested and expanded. The gel was imaged for red (OligodT-ATTO590) and DAPI (blue) fluorescence, which revealed both nuclear and cytoplasmic presence of captured mRNAs. (D-K) Comparison of transcriptome capture coverage in liver tissue across various spatial transcriptomics (ST) technologies. Six publicly available ST datasets (D-J) and the current Seq-Scope-X dataset (K) are shown to illustrate the spatial distribution of transcriptome capture coverage. The datasets include: Original ST (D) [25], 10x Visium (E) [26], Slide-Seq (v1; F) [8], xDBiT-Seq (G) [27], Seq-Scope^MISEQ^ [12] (H), Stereo-Seq (I) [28], Seq-Scope^NOVASEQ^ [15] (J), and the Seq-Scope-X dataset (K). The Seq-Scope-X approach combines Seq-Scope^NOVASEQ^ with tissue expansion to enhance spatial resolution. For standardized comparisons, datasets were binned into grids or hexagons, depending on data availability: Unmodified (Original ST, 10x Visium and xDBiT), 50 µm × 50 µm grids (Stereo-Seq), 10 µm × 10 µm grids (Seq-Scope^MISEQ^), 14 µm-sided hexagons (Seq-Scope^NOVASEQ^), and 5 µm-sided hexagons (adjusted for expansion; Seq-Scope-X). Area-normalized transcriptome capture efficiencies for each feature, bin or grid were visualized as spatial heat maps (D-K). The total distribution of transcriptome capture performance is summarized in Fig. 1L. White scale bars represent the original scale, while yellow scale bars indicate the scale adjusted to account for tissue expansion or computational dilation.

**Fig S2.**
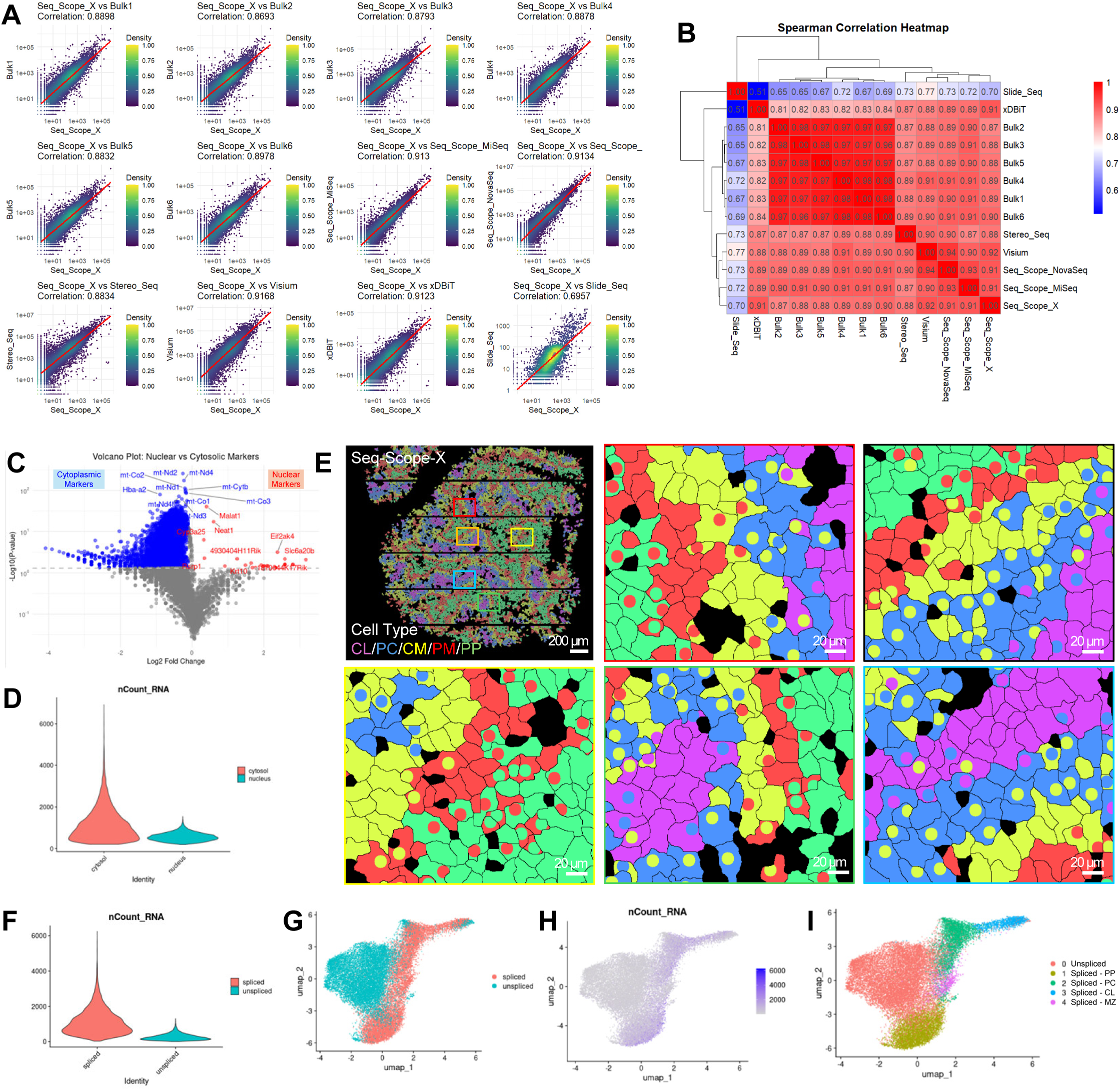
Characteristics of Seq-Scope-X Data from Mouse Liver. (A and B) Comparison of mouse liver transcriptomes from different spatial transcriptomics (ST) technologies and bulk RNA-seq. Datasets include six batches of bulk RNA-seq (Bulk1-6) [29], 10x Visium [26], Slide-Seq (v1) [8], xDBiT-Seq [27], Stereo-Seq [28], Seq-Scope^MISEQ^ [12], Seq-Scope^NOVASEQ^ [15], and the Seq-Scope-X datasets. All data were produced from young (2-4 month old) C57BL/6 male mice, except Slide-Seq (v1) data whose animal source was not identified. To facilitate comparisons, all ST datasets were aggregated into pseudo-bulk gene expression count tables. (A) Scatterplots show correlations between datasets, with each dot representing gene expression in two datasets. The red trendline indicates the correlation, and Spearman correlation coefficients (r) are displayed. (B) Heat map with dendrogram indicates the strong correlation (r>0.8) among all liver bulk RNA-seq and ST datasets except Slide-Seq (v1). The deviation of Slide-Seq data could be due to the biological difference due to different age, sex or strains. (C) A volcano plot shows genes differentially expressed between nuclear (red) and cytoplasmic (blue) compartments. As expected, nuclear-enriched non-coding RNAs, such as *Malat1* and *Neat1*, are enriched in nuclear segments, while mitochondrial genes are highly enriched in cytoplasmic segments. (D) A violin plot compares the number of transcripts in nuclear (red) and cytoplasmic (blue) segments, showing slightly fewer transcripts in nuclear segments. (E) Segments are color-coded based on cell type mapping results and visualized using the same color scheme as Fig. 2C and 2D. Regions not represented in Fig. 2C are included in these panels to provide additional context. (F-I) Spliced and unspliced transcriptome analysis based on whole-cell segments of Seq-Scope-X data. (F) A violin plot shows the number of spliced (red) and unspliced (green) transcripts in different whole cell segments, showing significantly fewer unspliced transcripts in whole cell segments. (G-I) UMAP visualizations of cell type clustering based on spliced and unspliced transcriptomes. (G) Data points are color-coded to distinguish spliced (red) and unspliced (green) data. (H) Data points are colored by the number of unique transcripts identified in each data. (I) Cell type clustering analysis highlights hepatocellular zonation clusters in the spliced dataset. Clusters 1-4 represent periportal (PP), pericentral (PC), centrilobular (CL) and midzone (MZ) hepatocytes, all corresponding to spliced data. In contrast, the unspliced data forms a single globular cluster (cluster 0), likely due to its sparse and biased transcriptome content. All scale bars indicate the scale adjusted to account for tissue expansion.

**Fig S3.**
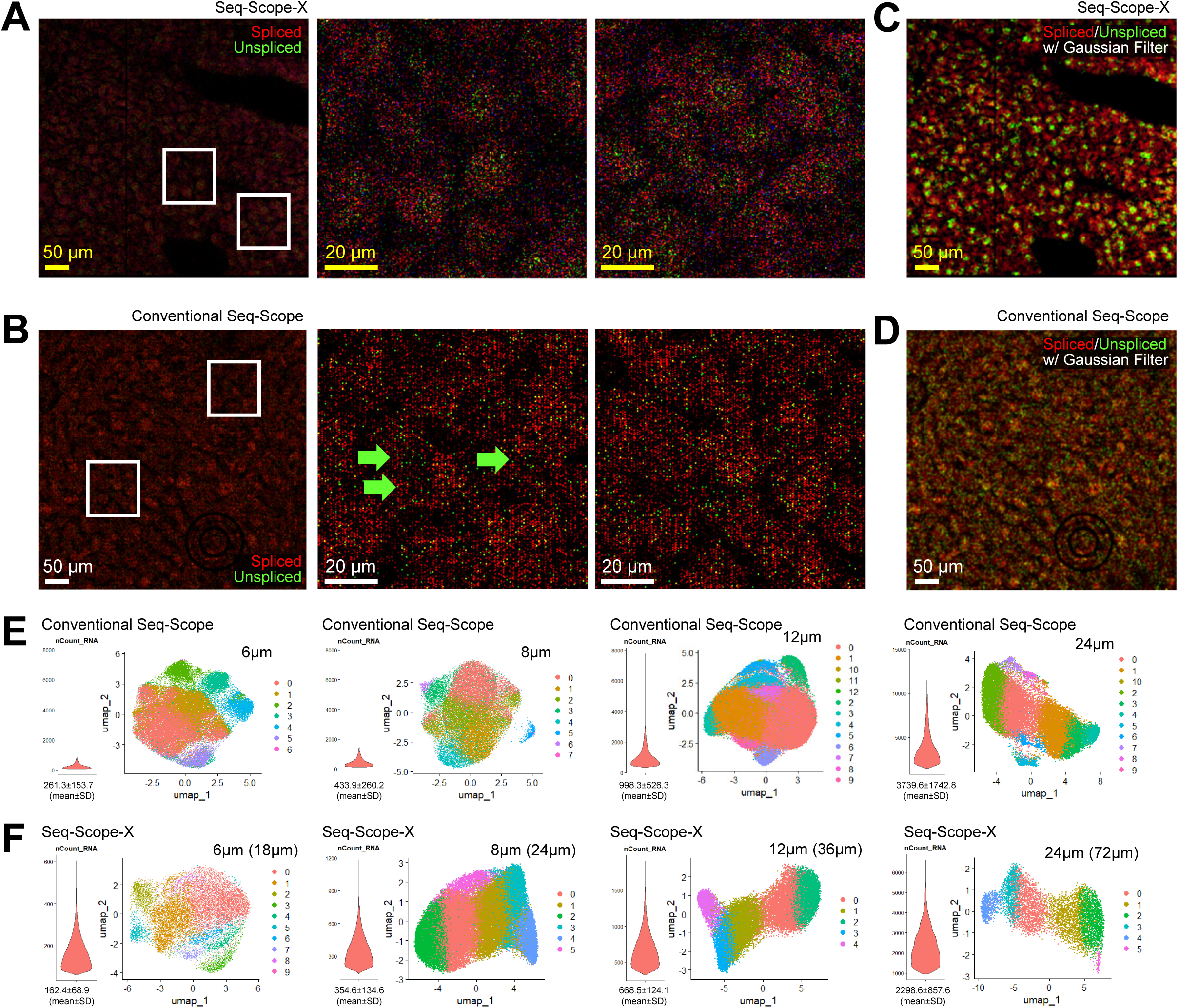
Comparative Spatial Performance of Seq-Scope-X and conventional Seq-Scope. (A and B) Spatial plotting of spliced (red) and unspliced (green) transcripts in Seq-Scope-X (A) and conventional Seq-Scope (B) data. In this comparison, the data are shown in the same effective magnification. Single cell and subcellular structure shown in (A) is not obvious in (B). Arrows highlight occasional detection of nuclei-like unspliced transcript clusters in conventional Seq-Scope data. (C and D) Digitally enhanced images of spliced (red) and unspliced (green) transcripts processed using Gaussian filtering modeling the same diffusion distance in Seq-Scope-X (C) and conventional Seq-Scope (D) data. (E and F) UMAP manifolds display cell type clustering using hexagons with specified flat-to-flat height (in µm) for conventional Seq-Scope (E) and Seq-Scope-X (F) liver datasets. Heights in parentheses (F) represent raw flat-to-flat dimensions, unadjusted for the expansion factor. Hexagons with a 24 μm height (unadjusted scale) correspond to the 14 μm-sided hexagons used in Fig. 3A and 3D. While the overall cluster geometry is similar between datasets, Seq-Scope-X demonstrates significantly improved spatial precision, as shown in Fig 3. White scale bars represent the original scale, while yellow scale bars indicate the scale adjusted to account for tissue expansion.

**Fig S4.**
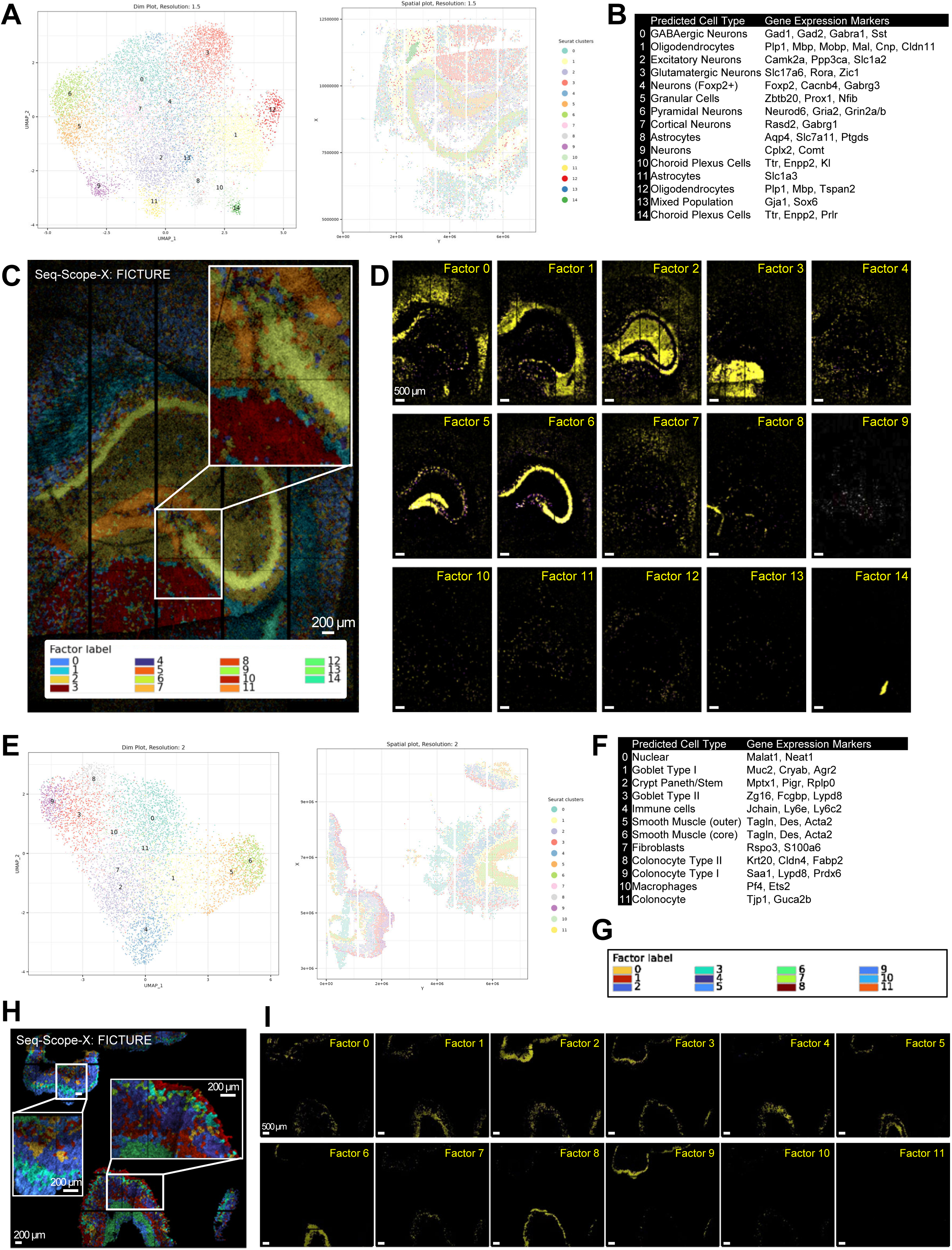
Detailed Tissue and Cell Type Insights from Seq-Scope-X Analysis. (A-I) Cell type clustering and spatial projection results of mouse brain (A-D) and colon (E-I) Seq-Scope-X datasets, described in Fig. 4. (A, B, E, and F) Cell type clustering results are shown as UMAP manifolds (A and E, left) and spatial maps (A and E, right) using 5 μm-sided hexagons (8 μm flat-to-flat height, reflecting 3X tissue expansion) from the Seq-Scope-X dataset. The same cell type-specific color code is applied for both plots. Cluster annotations and representative marker genes are summarized in a table for each dataset (B and F). The same clustering results were used for FICTURE projections (C, D, and G-I). (C, D, and G-I) FICTURE projections of the cell type clustering results in (A, B, E, and F) are presented in composite figures (C and H) with cluster-specific color codes (C, inset, and G) or as individual panels highlighting each factor separately. For the colon results (H and I), FICTURE output was spatially filtered using a mask to visualize only the expressions within the tissue boundary. All scale bars indicate the scale adjusted to account for tissue expansion.

**Fig S5.**
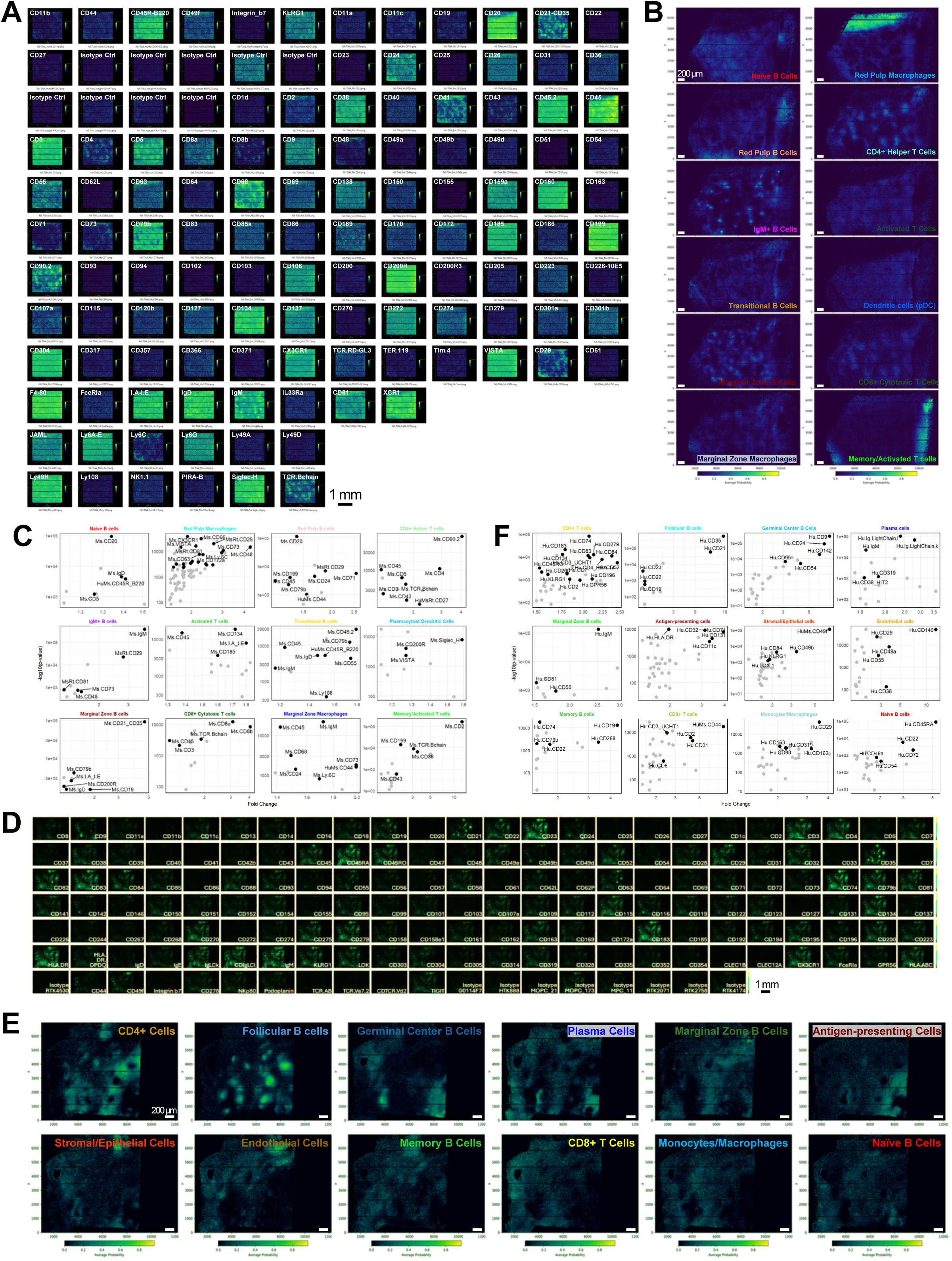
Spatial Proteome Analysis of Mouse Spleen and Human Tonsil Using Seq-Scope-X. (A-C) Seq-Scope-X Analysis of Spatial Proteome in Mouse Spleen. (A) Spatial expression plots of all 128 AbTags used for profiling the mouse spleen spatial proteome, as shown in Fig. 5B-5F. (B) Individual maps of all 12 LDA factors described in Fig. 5E. (C) Volcano plot of proteins highly expressed in pixels corresponding to each factor. Black dots represent proteins consistent with the annotated factor identity, while gray dots represent proteins that may not align with the annotated cell type but could reflect regional context. (D-F) Seq-Scope-X Analysis of Spatial Proteome in Human Tonsil. (D) Spatial expression plots of all 163 antibody tags used for profiling the human tonsil spatial proteome, as shown in Fig. 5G-5K. (E) Individual maps of all 12 LDA factors described in Fig. 5G, illustrating the positional probability of each factor. (F) Volcano plot of proteins highly expressed in pixels corresponding to each factor. Black dots represent proteins consistent with the annotated factor identity, while gray dots represent proteins that may not align with the annotated cell type but could reflect regional context. All scale bars indicate the scale adjusted to account for tissue expansion.

**Fig. S6.**
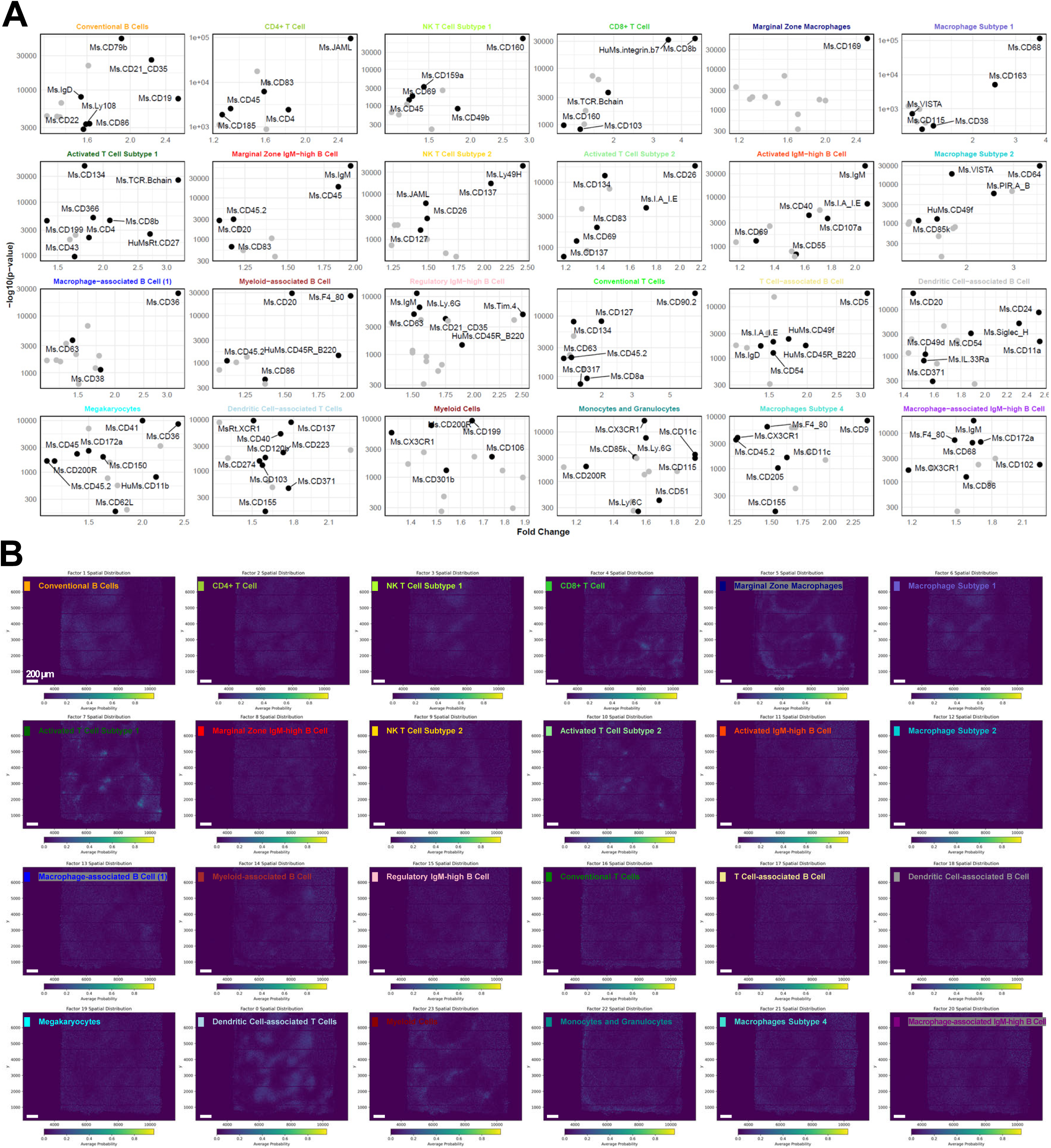
Analysis of the Mouse Spleen Spatial Proteome Profiled by DMAA-Enhanced Seq-Scope-X. (A) Volcano plot showing proteins highly expressed in pixels corresponding to each LDA factor described in Fig. 6. Black dots represent proteins consistent with the annotated factor identity, while gray dots indicate proteins that may not align with the annotated cell type but could reflect regional context. (B) Individual maps of all 24 LDA factors described in Fig. 6, demonstrating the positional probability of each factor within the histological space. All scale bars indicate the scale adjusted to account for tissue expansion.

